# Binary partitioning of human brain organization due to divergent human cytoskeletal evolution

**DOI:** 10.64898/2026.05.18.725959

**Authors:** Nicolás Rosas, Ivanna Ihnatovych, Jack A. Reeves, Eduardo Cortes Gomez, Ryu P. Dorn, Harneet Sandhu, Anna Szombathy, Niels Bergsland, Norbert Sule, Sarah Muldoon, Robert Zivadinov, Ralph H. B. Benedict, David A. Bennett, Jianmin Wang, Kinga Szigeti

## Abstract

*CHRFAM7A* is a biallelic uniquely human fusion gene present in 99.3% of humans. The direct and inverted alleles likely emerged independently in Africa and East Asia. We uncovered that the inverted allele regulates *ULK4* expression through genetic epistasis. The increase in long to short *ULK4* isoform ratio enhances α-tubulin acetylation leading to microtubule (MT) cytoskeleton phenotypes (cell body: microtubule rich projections; neurite: de-bundling; growth cone: fanning with MT invasion). The MT cytoskeleton gain of function enhances neuronal arborization and functional connectivity in the human brain. Considering that the previously reported direct allele leads to actin cytoskeleton gain of function, we propose that *CHRFAM7A* alleles represent binary human genetic background through divergent evolution of the cytoskeleton. In the human brain the two alleles represent distinct brain organization: the direct allele enhances microstructure as measured by diffusion tensor imaging while the inverted allele increases functional connectivity with increased small world propensity. The two types of brain organization likely represent different susceptibility to neuropsychiatric disease and may underlie the allelic disease associations.

**Graphical Abstract:** 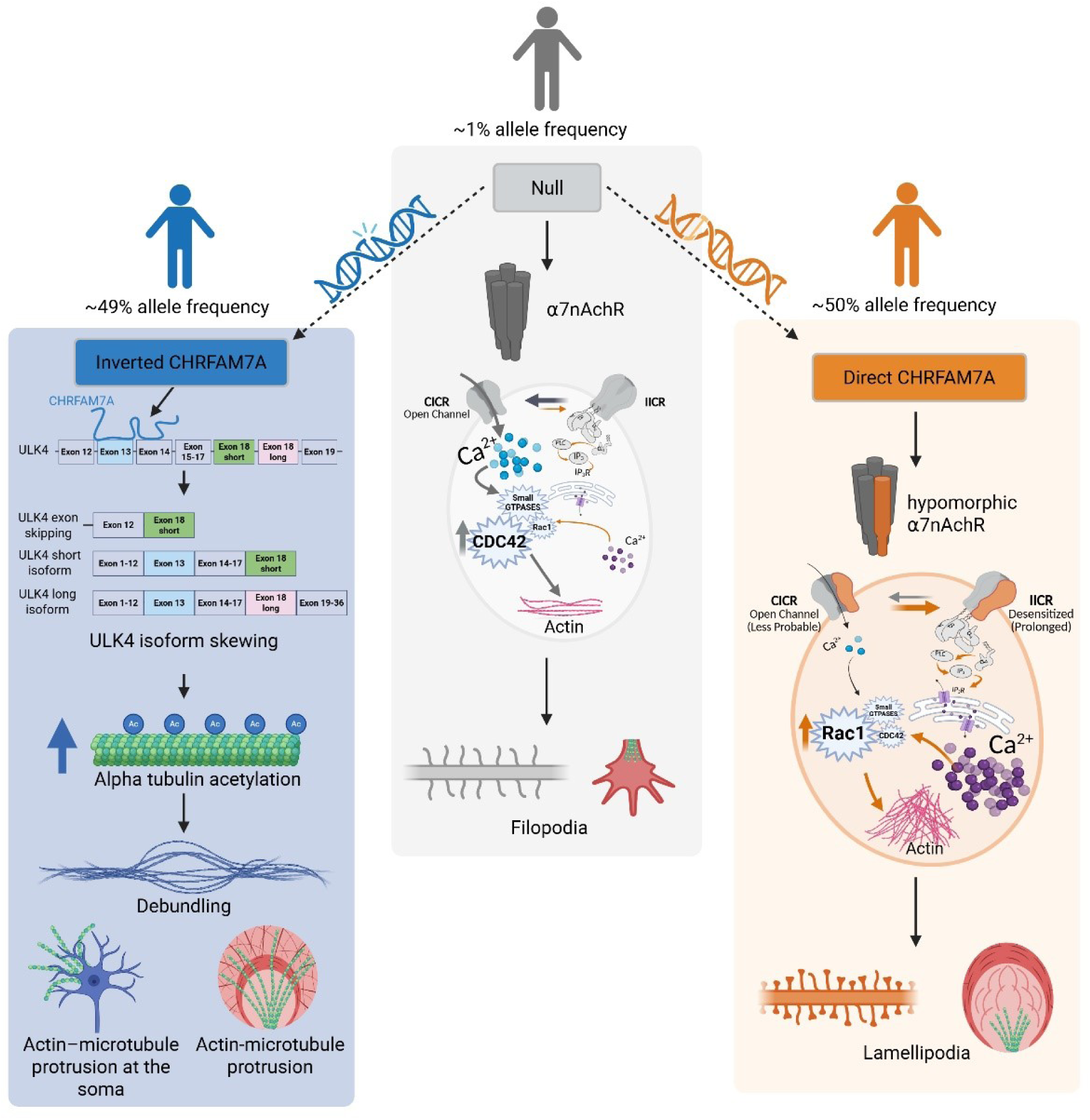

## Introduction

*CHRFAM7A* is a fusion gene between part of *CHRNA7* (exons 5-10) and part of *FAM7A (ULK4). CHRFAM7A* can be present in direct or inverted orientation (Fig. 1A) ^1,2^. Divergent human evolution of the *CHRFAM7A* direct and inverted alleles reflects a uniquely human pattern of structural variation shaped by duplication, inversion, and recombination events on chromosome 15 ^3^. The 1000 Genomes Project provides population-level evidence that these alleles segregate differently across human groups. In Africa the inverted allele frequency is 8% while in East Asian population it is 67% ^3^. The complexity of the rearrangements and difficulty deriving one from the other makes independent events more likely. Together, the 1000 Genomes data and structural analyses suggest that the two alleles represent distinct evolutionary trajectories within modern humans, shaped by both ancient rearrangements, selective pressure and ongoing population-specific variation. The functional characterization of the direct *CHRFAM7A* allele revealed an actin cytoskeleton gain of function affecting fundamental processes in the immune and nervous systems ^4,5^. The function of the inverted allele remained elusive.

**Figure 1:**
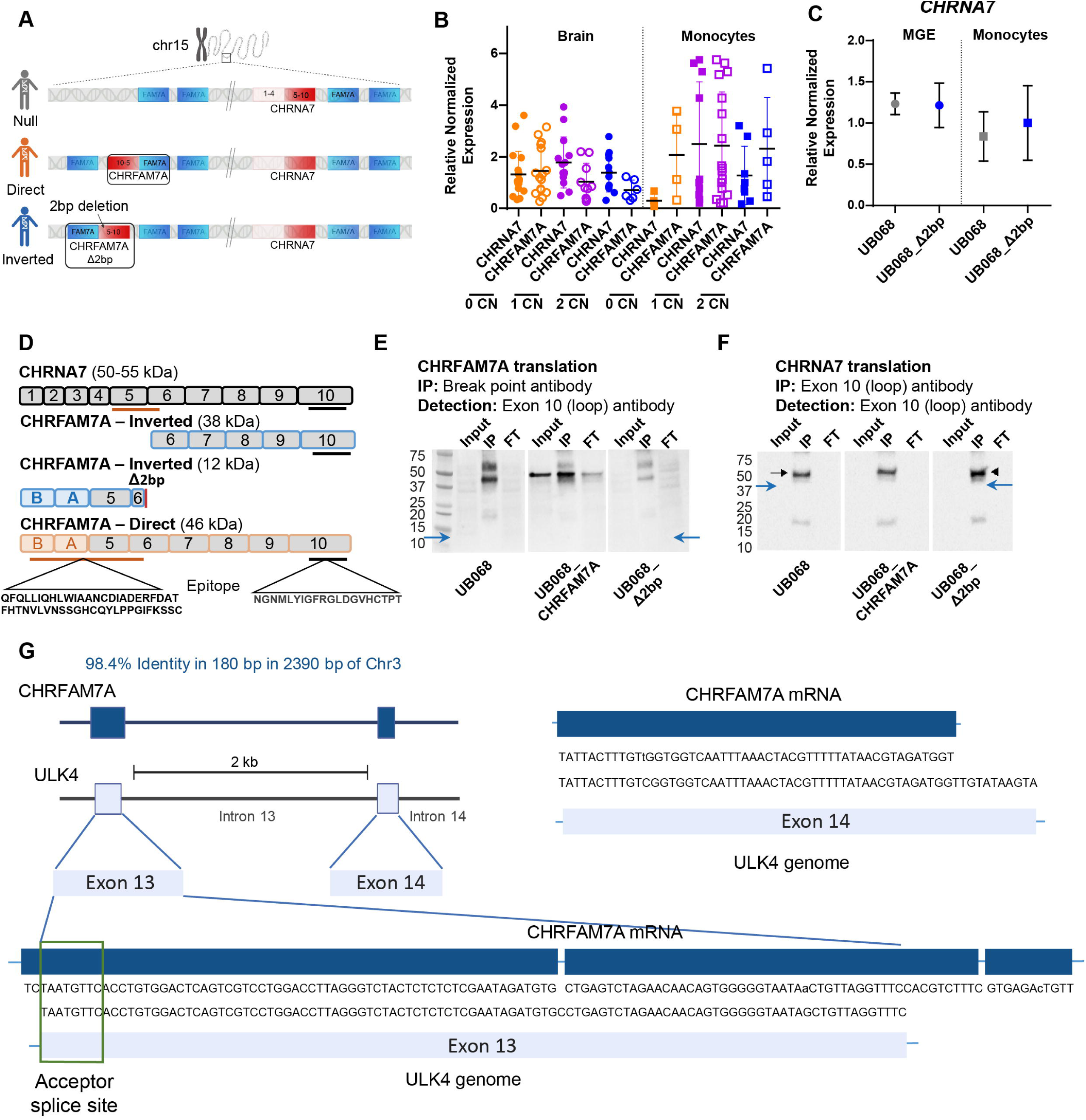
Inverted CHRFAM7A functions as a regulatory RNA. **A.** Schematic of chr15:30,44. The ancestral allele contains both the *CHRNA7* and *FAM7A*. Partial duplication and fusion of *CHRNA7* and *FAM7A* generates *CHRFAM7A.* **B.** *CHRFAM7A* and *CHRNA7* gene expression in post mortem human brain and primary human monocytes in the presence of 0 (orange), 1 (purple) and 2 (blue) copies of the inverted allele. **C**. Inverted *CHRFAM7A* does not affect *CHRNA7* expression in MGE progenitors and monocytes differentiated from the null (UB068) and the inverted (UB068_Δ2bp) isogenic lines. **D.** Schematic representation of the protein sequences and corresponding molecular weight predicted from the inverted CHRFAM7A sequence harboring the 2bp deletion. **E.** Western blots demonstrating absent CHRFAM7A bands in the null and inverted iPSC derived MGE progenitors contrasted to the direct CHRFAM7A expressed protein at 46 kDa. **F**. *CHRNA7* encoded α7 nAChR subunit is expressed in all 3 cell lines. Blue arrows show the predicted sizes of inverted allele proteins 38 kDa and 12 kDa. FT, flow through. **G.** Direct and inverted *CHRFAM7A* EST sequence mapped to the genome browser reveals a single hit exhibiting 99% identity to exon 13 and exon 16 of *ULK4*.

Both participants in the fusion event are important neuronal genes. *CHRNA7* codes the subunit of the α7 nAChR homopentamer. The α7 nAChR plays a role in cognition and memory and has been a leading druggable target for Alzheimer’s disease and schizophrenia, albeit hindered by a persistent translational gap ^1,6–12^. ULK4 belongs to the Unc-51-like kinase family, but unlike classical serine/threonine kinases, ULK4 lacks key catalytic residues and functions primarily as a scaffolding or regulatory protein rather than an active enzyme ^13,14^. During early neurodevelopment, alternative splicing tends to favor the short ULK4 transcript, which encodes a truncated protein lacking the extended C-terminal protein-protein interaction region ^15^. Alternative splicing is highly utilized in neuronal development, frequently affecting the cytoskeleton ^16^. As development progresses—particularly during neuronal differentiation and maturation—splicing shifts toward the full-length long isoform, producing the ∼1,275-amino-acid protein with its complete pseudokinase domain and extensive C-terminal modules ^15^. This transition will likely expand ULK4’s interaction capacity, enabling more complex signaling roles in neurite outgrowth and microtubule stability ^17^.

ULK4 microdeletion syndromes are associated with schizophrenia and KO mouse models deciphered a role of ULK4 in neuritogenesis, migration, branching and network development ^14,15,18^. ULK4 KO mice demonstrated anxiety like behavior associated with altered GABAergic differentiation ^19^.

It has been recently shown that the direct *CHRFAM7A* allele is translated and produces a hypomorphic α7 nAChR with functional consequences of modifying Ca^2+^ signaling, resulting in altered Rac1 activation and actin cytoskeleton gain of function ^4^. The inverted allele harbors a 2 bp deletion in exon 6 of the *CHRNA7* derived sequence leading to a frameshift mutation ^20,21^.

The equal allele frequencies of the direct and inverted alleles in the North American admixture suggest that the inverted allele underwent similar selective pressure as the direct allele, implying that the inverted allele is functional.

Transcription of the inverted allele specifically has not been reported. Of note, gene expression studies of *CHRFAM7A* detect RNA transcribed from both alleles. *CHRFAM7A* expression has been reported in various cell types and under normal and pathological conditions ^11,22,23^. Translation of the inverted allele has not been demonstrated. In silico analysis predicted three scenarios; the protein is not translated due to the distance between the Kozak consensus sequence from initiation ^21^; truncated peptide is translated due to an early stop codon; or truncated *CHRNA7* is translated from exons 5-10^11,24^. Functional studies in neuronal and mononuclear lineage derived from human isogenic iPSCs indicate that the inverted allele is a functional null from the α7 nAChR perspective ^2^. Mechanistically it has been unclear what drives the selection pressure for the inverted allele.

While physiological function is still unknown, there is a reasonable signal that inverted *CHRFAM7A* allele is a risk factor for psychiatric diseases ^25–29^. These associations are detected with a candidate gene approach due to genomic architectural challenges with the locus, including limited mappability and sparse marker coverage ^1^.

Taking together that translation of the inverted allele is unlikely and it does not affect α7 nAChR function, we hypothesize that the inverted allele functions at the RNA level. To decipher the function of the inverted *CHRFAM7A* allele we utilize post mortem human brain, human primary monocytes as an accessible tissue and MGE progenitors and monocytes/macrophages differentiated from the isogenic iPSC model ^30^.

## Results

### Inverted allele is transcribed in human brain and primary monocytes

Gene expression of *CHRFAM7A* has been reported; however the utilized assays were not allele specific due to challenges in capturing the breakpoint and the 2 bp deletion in the same assay (Fig 1A). Allelic *CHRFAM7A* gene expression was explored in genotyped post mortem human brain tissue (N=48, subset of Supp. Table 1), human primary monocytes and macrophages (N=43, Supp. Table 2) in homo- and hemizygous direct and inverted genotypes and in heterozygotes. In samples that are homo- and hemizygous for the inverted allele, *CHRFAM7A* expression can only originate from the inverted allele, indicating that the inverted allele is transcribed (Fig. 1B).

### The inverted allele is not translated

Detection of CHRFAM7A at the protein level poses additional challenges. In the CHRFAM7A fusion protein, CHRNA7 is the C-terminal and ULK4 fragment is the N-terminal of the proposed protein products (Fig. 1D). The unique N-terminal aminoacid (AA) sequence does not have any commercially available antibodies. Antibodies targeting the breakpoint would theoretically be specific for CHRFAM7A; the C-terminal antibodies detect both CHRNA7 and CHRFAM7A (Fig. 1D). Protein translation has been demonstrated from the direct allele by us and others and serves as a positive control ^31–33^. The predicted protein product from the inverted allele that harbors the C-terminal AA sequence should be detectable by commercially available antibodies and can be distinguished from CHRNA7 by size on Western blots (Fig. 1D and Supplementary Fig. 1).

We performed a series of IP and Western blot experiments using our isogenic cell lines and we did not detect CHRFAM7A protein in the inverted iPSC line; direct iPSC line demonstrates the expected size and the band is absent from the null iPSC line (Fig. 1E). All three iPSC lines express CHRNA7 (Fig. 1F). We previously reported α7 nAChR functional studies including neurophysiology (patch clamp) and Fluoro4 live Ca^2+^ imaging where we found that the inverted allele does not affect the α7 nAChR, which are also indicative of lack of functional protein translation ^2^. We proceeded to study whether the inverted sequence may affect any of the fusion partners.

### Inverted CHRFAM7A functions through genetic epistasis on ULK4

*CHRNA7* expression level was not affected by the inverted allele in human brain or primary monocytes (Fig. 1B), nor in the isogenic iPSC derived MGE progenitors (Fig. 1C). Mapping of *CHRFAM7A* mRNA to the human genome sequence (T2T CHM13v2.0/hs1) revealed a match on chromosome 3 corresponding to *ULK4* exons 13 and 14. The alignment resulted in a 180 bp region with a 98.4% identity. Exon 13 harbors 99.3% identity spanning 127 bp within this region, with CHRFAM7A sequence overhanging both proxymal and distal exon 13 bounderies. Exon 14 has 98% identity over 50 bp within the region. The high identity score affecting adjoining exons and splice sites raise the possibility of genetic epistasis between *CHRFAM7A* and *ULK4* (Fig. 1G).

We utilized human brain RNAseq data (N=205 from ROSMAP, Supplementary Table 1) to explore correlation between *ULK4* transcripts in samples harboring 0, 1 and 2 copies of *CHRFAM7A* inverted allele. We detected three transcripts in more than 50% of the samples, long, short and mini (Fig. 2A). Long and short are the two translated *ULK4* transcripts, and they harbor exons 1-18 (short) and 1-37 (long) coding the short and long isoform of ULK4, respectively. Exon 18 is different in long and short isoform through alternative splicing (Fig. 2G). Both short and long isoforms were previously detected in the human brain ^15^. In the studied human brain cohort, the long isoform expression trended with inverted allele copy number (Fig. 2A). The isogenic iPSC model-derived MGE progenitors demonstrated increased long to short ULK4 ratio in the inverted line compared to null (Fig. 2B). We performed Spearman correlation on gene expression levels of the long and short transcripts (RNAseq data, N=205 from ROSMAP) and found decreasing correlation coefficient between short and long as a function of inverted allele copy number (Fig. 2C). To validate and refine these findings we performed long and short *ULK4* transcript specific qPCRs (based on different exon 18 sequence) on a subset of ROSMAP post mortem human brain (N=48) RNA (Fig. 2D) and in human primary monocytes (46 donors) (Fig. 2E). In both human tissues correlation between short and long isoforms decreased with increasing inverted allele copy number. These data suggest that the inverted *CHRFAM7A* allele may exert genetic epistasis on *ULK4* resulting in increased long to short *ULK4* isoform ratio.

**Figure 2:**
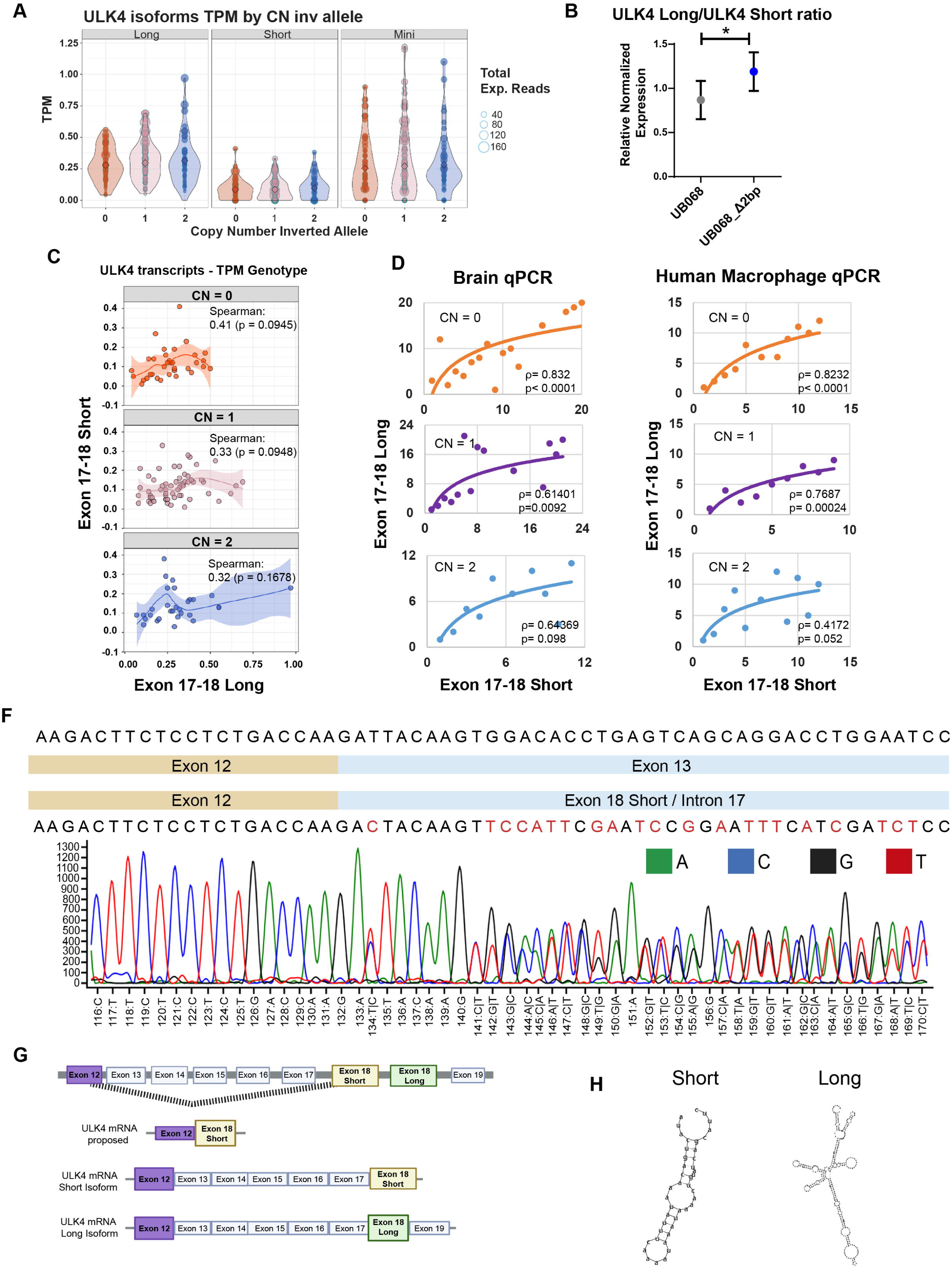
Inverted CHRFAM7A modifies ULK4 isoform ratio through genetic epistasis. **A.** *ULK4* isoforms (long, short and mini) captured with RNAseq in 100 human post mortem brain tissue (ROSMAP) is depicted in the presence of 0 (orange), 1 (purple) and 2 (blue) copies of the inverted allele. Transcripts per million is depicted on the Y axis, inverted allele copy number on the X axis. Size of circles are proportionate to the number of experimental reads. Long *ULK4* isoform reads demonstrate an inverted allele dosage effect. **B.** Increased long to short *ULK4* ratio is present in isogenic inverted UB068_Δ2bp MGE progenitors iPSC line. Data are means; *p < 0.05 UB068 Short vs UB068_Δ2bp Short and UB068 Long vs UB068_Δ2bp Long using two-sided unpaired t-test (n=5). **C.** Correlation between the *ULK4* isoforms in the presence of 0 (orange), 1 (purple) and 2 (blue) copies of the inverted allele: the short and long isoform correlation demonstrates an inverted allele dosage effect, less correlation with increasing number of inverted copy number. **D.** *ULK4* short and long isoforms demonstrate decreasing correlation with increased inverted *CHRFAM7A* copy number in human post mortem brain tissue (N=48) by qPCR. Exon-exon boundary specific assays designed to detect Short (exon17-exon18short) and long (exon17-exon18long) isoforms selectively. Statistical analysis by Spearman rank correlation. **E.** *ULK4* short and long isoforms demonstrate decreasing correlation with increased inverted *CHRFAM7A* copy number in human macrophages. **F.** Sanger sequencing chromatogram detects the exon 12-18sh alternatively spliced variant in the inverted MGE progenitors. **G.** Schematic of the proposed genetic epistasis between *CHRFAM7A* and *ULK4*. **H.** Predicted secondary RNA structure of 3’ end of short and long *ULK4* isoforms using UCSC database.

As the *CHRFAM7A* sequence maps to exon 13 and 14, we expected that its presence might alter splicing in this region of the *ULK4* transcript. To test this, we performed long-range RT-PCR using primers spanning exon 10 and exon 18short to amplify the short isoform, and exon 10 to exon 19 to amplify the long isoform exclusively. Sanger sequencing revealed an alternatively spliced product joining exon 12 to exon18short (Fig. 2F) in the inverted isogenic line, indicating exon skipping in short ULK4 isoform (Fig. 2G). Although exons 13 and 14 are also included in the long isoform, the long transcript has complex predicted secondary structure at the 3’ end of the RNA (Fig. 2H). The extended and branched conformation of the long RNA may promote more complex folding, potentially altering the epistatic relation between CHRMFA7A and ULK4. The splice acceptor site sequence of the skipped exon 13 (ACTACAAG) is identical to the splice acceptor site of exon 18sh except for a single base (ATTACAAG) which may account for the preference of exon 18sh in the splicing process. The conundrum of the detected skewing towards the long isoform may be accounted for by regulatory mechanism activated by lower short isoform expression driving *ULK4* gene expression. The shift toward increased long-isoform abundance may be explained by a regulatory mechanism triggered by the reduced expression of short isoform skewing the isoform balance in the inverted line. Western blot of UB068 and UB068_Δ2bp demonstrated the presence of both the long (150 kDa) and the short (40 kDa) ULK4 isoforms (Fig. 3A) in iPSC derived MGE progenitors (Fig. 3B); and the increased long to short isoform ratio in the inverted line compared to null (Fig. 3C). ICC for ULK4 detected more peripheral distribution of ULK4 in the inverted isogenic MGE progenitors compared to null (Fig. 3D), showing an increased cytoplasmic/membrane to nuclear ratio (Fig. 3E). Specificity of ULK4 signal detected by Western blot and ICC was confirmed by using an antibody specific blocking peptide (Supplementary Fig. 2).

**Figure 3:**
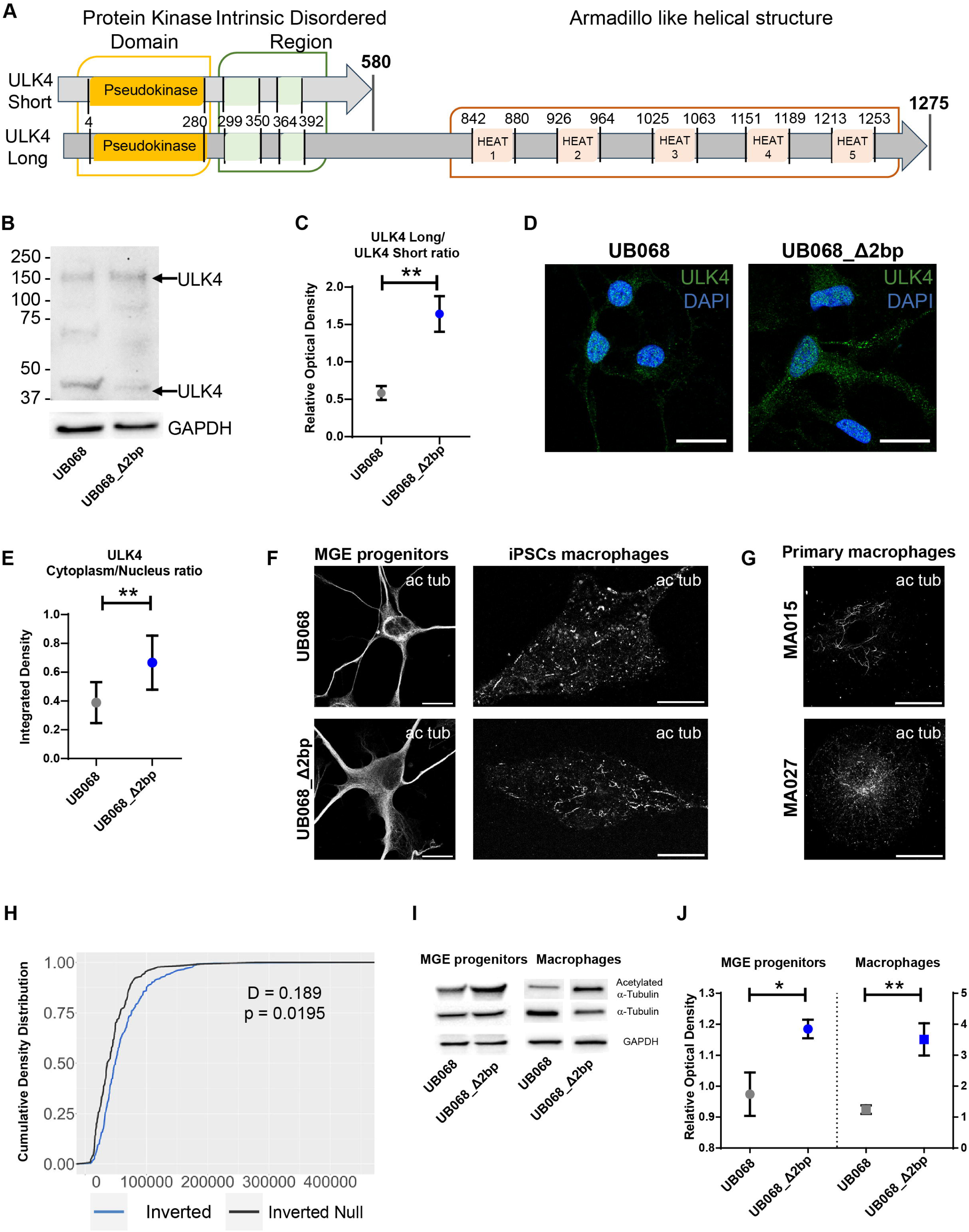
Increased long to short ULK4 isoform ratio facilitates a-tubulin acetylation. **A.** Schematic representation of the ULK4 isoforms. **B.** Western blot for ULK 4 demonstrates ULK4 long (150 kDa) and ULK4 short (40 kDa) detection in the isogenic UB068 and UB068_Δ2bp MGE progenitors. **C.** Densitometric analysis of ULK4 150 kDa and ULK4 40 kDa expression levels in the isogenic MGE progenitors shows an increased ULK4 long/ULK4 short ratio in the inverted MGE progenitors compared to the null. Data are presented as mean ± SD. **p < 0.01 using two-sided unpaired t-test (n=3). **D.** Representative images of isogenic MGE progenitors ICC for ULK4 (green) and DAPI (blue). Scale bar, 10 µm. **E.** Fluorescence density analysis of ULK4 levels between nucleus and cytoplasm in isogenic MGE progenitors. Data are presented as mean ± SD; **p < 0.01 using two-sided unpaired t-test (n=3). **F.** Representative confocal images of acetylated α-tubulin staining in isogenic null and inverted MGE progenitors (first column), and isogenic macrophages (second column). **G.** Representative confocal images of acetylated α-tubulin in the primary human macrophages with different copies of the inverted alleles. Scale bar, 10 µm. **H.** Cumulative density curves demonstrate higher level of α-tubulin acetylation in the inverted allele carriers compared to the inverted allele null. Two sample Kholmogorov-Smirnov statistics. **I-J.** Representative Western blot and densitometry analysis demonstrating increased acetylated to total α-tubulin ratio in the isogenic inverted MGE progenitors and macrophages compared to null. Data are presented as mean ± SD; UB068 vs UB068_Δ2bp monocytes *p < 0.05; UB068 vs UB068_Δ2bp MGE progenitors **p < 0.01 using two-sided unpaired t-test (n=3).

### Increased long to short ULK4 isoform ratio facilitates α-tubulin acetylation

As functional studies and mouse models implicate ULK4 in α-tubulin acetylation ^15,17,19^, next we explored whether the presence of the inverted allele is related to α-tubulin acetylation in iPSC derived MGE progenitors and macrophages, and human primary macrophages (N=20). Immunofluorescent microscopy for acetylated α-tubulin demonstrated increased peripheral acetylation in MGE progenitors differentiated from UB068_Δ2bp (Fig. 3F) coinciding with ULK4 peripheral distribution (Fig. 3G) and longer acetylated α-tubulin units in iPSC derived macrophages (Fig 3H). Human primary macrophages showed increased α-tubulin acetylation in the presence of inverted *CHRFAM7A* (Fig. 3G). Increased acetylated α-tubulin was detected by Western blot in inverted UB068_Δ2bp isogenic MGE progenitors and macrophages compared to UB068 null cells (Fig. 3I and J).

### Inverted allele leads to ULK4 associated microtubule cytoskeleton phenotypes

MGE neuronal progenitors labeled for acetylated α-tubulin showed marked differences in morphology affecting the whole neuronal structure including the cell body, neurites and growth cone (GC). In the soma, tubulin fibers of UB068_Δ2bp MGE progenitors splay and are less compact compared to null (Fig. 4A, first column). Actin and α-tubulin co-staining in isogenic progenitors revealed that these correspond to actin–microtubule protrusive structures at the soma (Fig. 4A, second column). These structures are characterized by being located solely in the cell body and having a lamellipodia conformation invaded by microtubules. In the neurites, tubulin fibers exhibit increased dynamic debundling (Supplementary video 2) with increased actin veil compared to null (Fig. 4A, third column). The GCs of inverted UB068_Δ2bp isogenic MGE progenitors have lamellipodia conformation with MT invasion which contrasts with null GCs harboring filopodia protrusions with limited microtubule invasion (Fig. 4A, fourth column).

**Figure 4:**
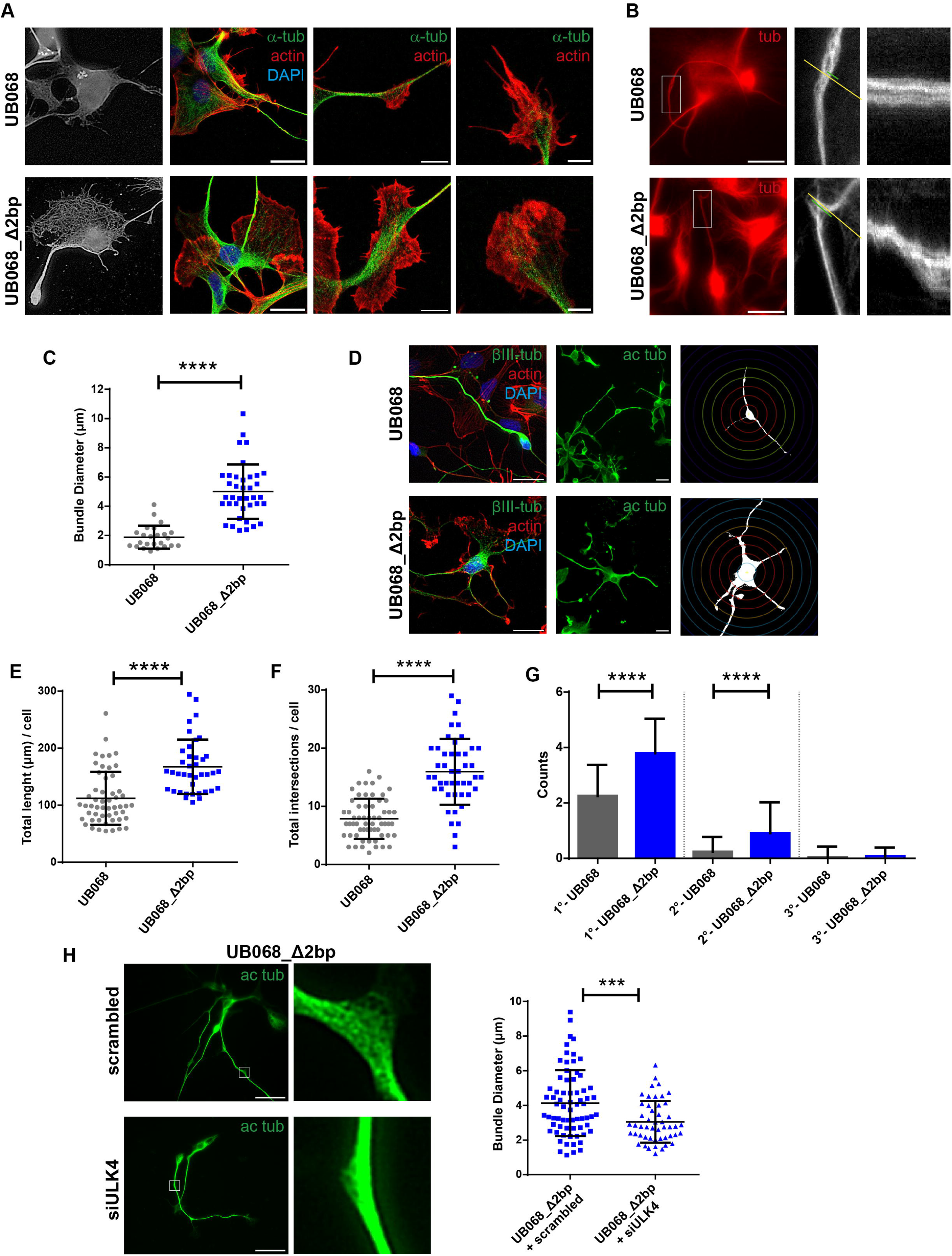
inverted allele is related to acetylated tubulin and actin phenotypes in isogenic MGE progenitors. **A.** Representative regions of UB068 and UB068_Δ2bp MGE progenitors. First column, cell body showing organization of microtubules stained with acetylated α-tubulin. Second column, cell body stained for α-tubulin (green), actin (red) and nuclei (blue). Third column, representative microtubule bundle region stained for α-tubulin (green), actin (red) and nuclei (blue). Fourth column, representative isogenic growth cones stained for α-tubulin (green) and actin (red). **B.** First column, representative images of live UB068 and UB068_Δ2bp MGE progenitors stained with SPY555-tubulin and the respective ROIs displaying enlarged axons segments. Second column, enlarged axons with green segments for measured bundle diameter and yellow for line used for kymograph. Third column, kymograph representing the dynamic movement of axons in a straight line during 304 seconds time course. Scale bar, 20 µm. **C.** Diameter of the tubulin bundle measured in a perpendicular line. Data are means ± SD; UB068 vs UB068_Δ2bp ****p < 0.0001 was assessed using unpaired two-tailed Mann-Whitney U test. **D.** First column, representative images of UB068_Δ2bp MGE progenitors stained with βIII-tubulin (green), actin (red) and DAPI (blue). Second column, UB068 and UB068_Δ2bp MGE progenitors stained for acetylated α-tubulin (first row) and Sholl analysis using Neuroanatomy plugin for measuring total intersections per cell (third row). Scale bar, 40 µm. **E.** Mean total length of neurite per cell calculated. Data are means ± SD; UB068 vs UB068_Δ2bp p**** < 0.0001 using unpaired two-tailed Mann-Whitney U test. **F.** Mean total intersection per cell individual cell calculated with Sholl analysis. Data are means ± SD; UB068 vs UB068_Δ2bp p**** < 0.0001 using unpaired two-tailed t-test. **G.** Type of process classification for primary neurite/axons, secondary neurite and tertiary neurite. Data are means + SD; primary and secondary neurites: UB068 vs UB068_Δ2bp p**** < 0.0001 using unpaired two-tailed Mann-Whitney U test. **H.** ROIs displaying microtubule bundles of UB068_Δ2bp MGE progenitors transfected with 100nM of siULK4 or scrambled-RNA for 96h stained with acetylated α-tubulin. Diameter of the tubulin bundle measured in transfected MGE progenitors. Scale bar, 40 µm. Data are means ± SD; UB068_Δ2bp scrambled vs siULK4 ***p < 0.001 from 2 independent experiments using unpaired two-sided t-test.

To gain further insights we performed Spy-555 live tubulin imaging in MGE progenitors. Neurites of UB068_Δ2bp MGE progenitors demonstrated areas of marked dynamic bending with debundling of tubulin fibers (Fig. 4B). Quantification of the width of microtubule bundles confirmed increased splaying in UB068_Δ2bp compared to UB068 (Fig. 4C). Focal debundling of parallel microtubule arrays has been reported to drive collateral axon branching ^34^.

Single cell Scholl analysis revealed increased total length of processes (Fig. 4D and E) and higher arborization (Fig. 4D and F) in UB068_Δ2bp progenitors. UB068_Δ2bp cells exhibited an increase in number of both primary (originating from soma) and secondary neurites. Tertiary neurites were rarely observed consistent with the early stage of differentiation (Fig. 4G). The increased number of primary neurites results in multipolarization in the presence of the inverted *CHRFAM7A* allele (Fig. 4G). In our differentiation protocol^30^ the majority of MGE progenitors differentiate into GABAergic neurons suggesting a role of inverted *CHRFAM7A* in enhancing GABA interneuron arborization and multipolarity.

To demonstrate ULK4 dependence, we used siRNA knockdown of ULK4 in the inverted MGE progenitors (Supplementary Fig. 3). Acetylated α-tubulin ICC was performed and debundling width was quantified. ULK4 knockdown eliminated the inverted allele inferred higher debundling in the inverted MGE progenitors (Fig. 4H).

### Inverted allele leads to enhanced locomotion

UB068_Δ2bp MGE progenitors show locomotion driven by the microtubule protusive structure where the direction of movement is generally directed toward the side of protrusion (Fig. 5A). Live imaging utilizing Life actin-RFP molecular probe reveals increased exploratory movement of neurites (quantified as distance covered, Fig. 5B and C) in UB068_Δ2bp progenitors compared to null (UB068). GC morphology is also more dynamic in UB068_Δ2bp progenitors and it is reversed by ULK4 siRNA KD, as illustrated by the temporal layering diagrams of the countour of the GC (Fig. 5D). These data suggest an increase in exploratory movement in inverted MGE progenitors which has implications for neurodevelopment, a prominent ULK4 phenotype ^17^.

**Figure 5:**
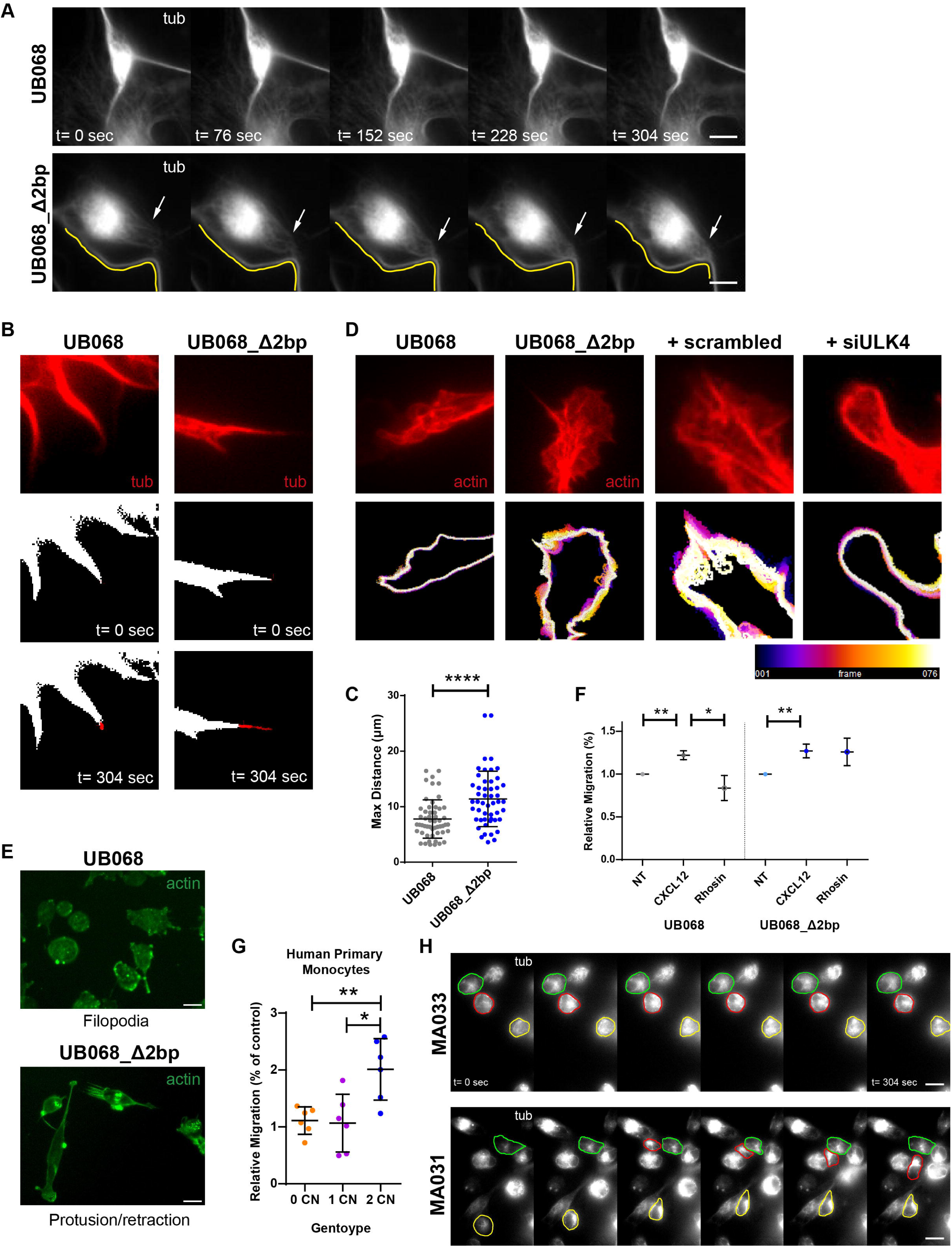
Inverted allele leads to increased cellular dynamics. **A.** Montage showing increased motility in UB068_Δ2bp MGE progenitors labeled by SPY555 tubulin molecular probe during a 304 sec time lapse. Arrows show regions of tubulin filament movement over an axon outlined in yellow. The center of the cell is circled in red. Scale bar, 10 µm. **B.** First row, ROIs of 10 µm^2^ of live images showing neurites of MGE progenitor transfected with Life actin-RFP. Second and third row, the resulting binary image at start (t= 0 sec) and end (t= 304 sec) of analysis. **C.** Max distance traveled of the tips of the neurites quantified using MTrackJ plugin from the ROIs (76 frames, total time= 304seg). Data are represented as mean ± SD from 4 independent experiments, unpaired two sided t-student test. **D.** First row, ROIs of 15 µm^2^ for growth cones and the resulting outline of the surface of the growth cone at start of image (t= 0 sec) and the end (t= 304 sec). Bottom row is the temporal color code for growth cones time course. **E.** Isogenic monocytes stained for actin displaying filopodia in UB068 and retraction and protrusion in UB068_Δ2bp. **F.** Migration as percentage of UB068 and UB068_Δ2bp isogenic monocytes treated with CXCL12 and Rhosin relative to non-treated control. Data are means ± SD; UB068 non-treated vs UB068 CXCL12 p** < 0.01; UB068 CXCL12 vs UB068 Rhosin p* < 0.05; UB068_Δ2bp non-treated vs UB068_Δ2bp CXCL12 p** < 0.01. Unpaired two-tailed t-test. **G**. Migration of human primary monocytes after CXCL12 exposure relative to control as percentage in the presence of 0 (orange), 1 (purple) and 2 (blue) copies of the inverted allele. Data are means ±SD; 1 copy vs 0 copy p** < 0.01; 1 copy vs 2 copy p* < 0.05 using unpaired two-tailed t-test. **H.** Montage showing increased motility in MA033 (0 copy) and MA031 (2 copy) human monocytes and stained with tubulin during a 304 sec time lapse. Color circles highlight the movement of individual monocytes.

As monocytes express *CHRFAM7A*, and microtubule cytoskeleton is key in cellular motility ^35^, we set out to investigate whether the MT gain of function is cell-type independent. UB068_Δ2bp isogenic monocytes demostrated increased polarization and protrusion-retraction phenotype by fluorescent phalloidin ICC (Fig. 5E) and increased migration compared to null (UB068) (Fig. 5F). Pharmacological modulation with RhoA small GTPase inhibitor showed that while migration is RhoA dependent in null (UB068) (reduced by RhoA antagonist Rhosin) (Fig. 5F), in UB068_Δ2bp isogenic monocytes it is RhoA independent (Fig. 5F). Human primary monocytes (N=18) demonstrated that inverted monocytes migrate more efficiently through transwell than direct or heterozygous counterparts (Fig. 5G). In primary human monocytes, the migration phenotype associated the inverted allele is characterized by MT polarization and cellular shape remodeling, captured in real time with the Spy555 MT molecular probe (Fig. 5H and Supplementary video 1).

### Brain functional connectivity increases as a dosage of the inverted allele

We designed a clinical-structural-functional study to characterize the inverted allele mediated brain phenotype using neuropsychological testing, structural MRI (diffusion tensor imaging; DTI) and functional MRI (CEG-MS, N=46). Allele specific dual genotyping identified 15 (33%) non-carriers, 22 (48%) 1-copy and 9 (20%) two-copy inverted allele participants. Genotyping was performed after the assessments. Demographic characteristics including age, sex, level of education is comparable between the groups (Supp. Table 3). We utilized the Baltimore Longitudinal Study on Aging (BLSA) autopsy cohort (N=96) baseline personality assessments that overlapped with our battery to enhance power. Demographics and genotype distribution for the BLSA cohort are depicted in Supp. Table 4.

In the clinical study comparison to the ancestral allele is not feasible as the null individuals are extremely rare (0.9% of the population). However, as the two *CHRFAM7A* alleles represent unrelated molecular mechanisms we can use the homozygous direct group as null for the inverted allele to explore inverted allele effect.

Processing speed, verbal learning and memory, visual memory, motor function, and depression and anxiety did not differ as a function of the inverted allele dosage (Supp. Table 5 and 5). Relative weakness in visual-spatial learning was detected (B = -5.578 CI=[-10.307, 0.850]; p = 0.021, uncorrected) in 2 copy inverted allele carriers (Supp. Table 5). In contrast, strength in openness personality trait in the NEO-FFI was positively associated with inverted allele dosage (B = 7.153 CI = [4.191, 10.116]; p = 0.005, Bonferroni corrected) (Supp. Table 6).

Volumetric analysis demonstrated similar whole brain, grey matter, white matter and ventricular volumes irrespective of inverted allele copy number (Supp. Table 7). DTI identified a combination of lower white matter fractional anisotrophy (FA) along with increased axial diffusivity (AD), medial diffusivity (MD), and radial diffusivity (RD) (relative weakness in microstructure) in 2 copy inverted participants with an assymetry lateralizing to the right side (Fig. 6D). Tract-Based Spatial Statistics with significant clusters of p < 0.05 resulted in peak MNI localizing to the right inferior fronto-occipital fasciculus, right superior longitudinal fasciculus and right corticospinal tract (Supp. Table 8). These pathways build a functional circuit that integrates visual sensory input with attentional control and motor output resulting in more flexible adaptation to complex spatial demands that support more effective interaction with dynamic environments. This relative weakness of the inverted allele is in fact the strength of the direct allele (0 copy inverted).

**Figure 6:**
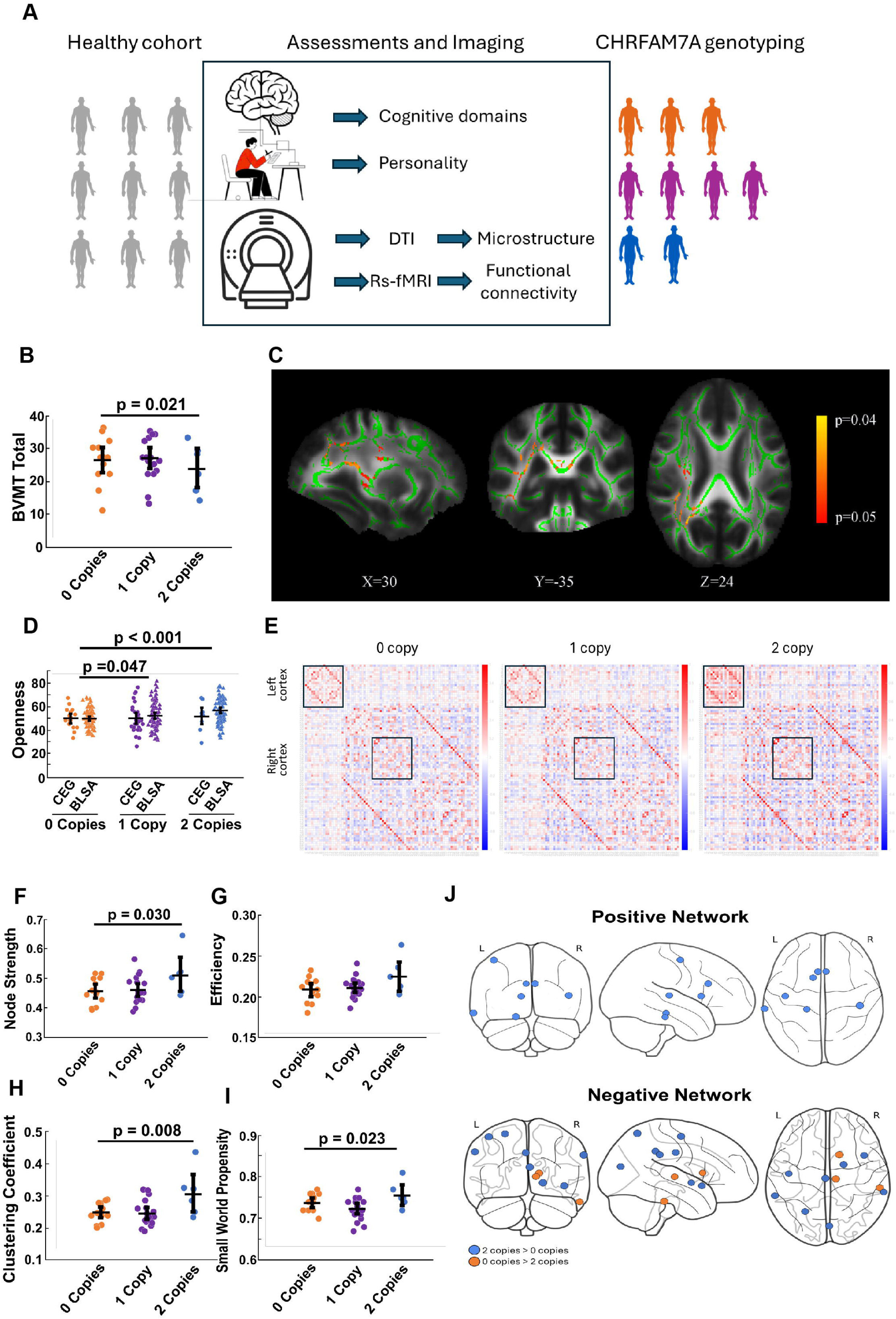
Cognitive performance, personality, and structural and functional human brain measures as a function of inverted allele dosage. **A.** Study design: healthy controls (N=46) underwent cogitive assessment, NEO-FFI, structural and functional MRI. Dual allele specific genotyping was performed after the assessement. **B.** Visual-spatial learning (BVMT) was lower in inverted homozygous individuals compared to inverted null (direct homozygotes, p = 0.021). **C.** Tract-based spatial statistics with the mean fractional anisotropy (FA) image shown in greyscale, the mean FA skeleton in green, and voxels with family-wise error p < 0.05 shown in red-yellow. Significant voxels were detected using threshold-free cluster enhancement in conjunction with non-parametric combination of FA, axial diffusivity (AD), mean diffusivity, (MD) and radial diffusivity (RD) images. Significant voxels indicate areas where individuals with two inverted alleles had a combination of decreased FA along with increased AD, MD, and RD compared to those with no inverted alleles. Peak MNI localizes to the right inferior fronto-occipital fasciculus, right superior longitudinal fasciculus and right corticospinal tract. **D.** Openness personality trait (NEOFF1) was higher in inverted homozygous individuals (p < 0.001). **E.** Heatmap of Pearson correlation coefficients between brain regions (numeric coding for regions are in supp. information) detected by functional MRI reveal higher connectivity in inverted brain lateralizing to the Left. Graph theory analysis revealed average higher node strength (**F**, p = 0.030), global efficiency (**G**), average clustering coefficient (**H**, p = 0.008), and small-world propensity (**I**, p = 0.023) in the inverted brain. **J.** Positive-network node strengths that are greater in inverted homozygotes are depicted over glass brain model with localization to thesemantic-executive regions, with predominance on the Left side. Uncorrected p-values.

In contrast, the resting state functional MRI revealed that 2 copy inverted allele carriers demonstrated higher greater overall average node strength (B = 0.057 CI=[0.006, 0.108]) and small world propensity (B = 0.024 CI=[0.003, 0.045]) in the positive-correlation network (Fig. 6F-I and Supp. Table 9). In the negative-correlation network, global efficiency was increased (B = 0.020 CI=[0.002, 0.038]) in 2 copy inverted allele carriers (Supp. Table 9). Average clustering coefficient was associated with inverted allele dosage in both the positive- (B = 0.060 CI=[0.016, 0.105]) (Fig. 6H and Supp. Table 9) and negative-correlation networks (B = 0.042 CI=[0.003, 0.080]) (Supp. Table 9). Regions where inverted allele copy number is associated with positive and negative node strength are depicted in Fig. 6J and Supplementary Table 10. The constellation of regions identified reflects a neural architecture characterized by unusually strong integration across cognitive-control, language, memory, sensory, and motor systems. Enhanced connectivity in the anterior cingulate cortex, caudate nucleus, and lateral temporal regions supports efficient conflict monitoring, goal-directed behavior, and controlled semantic processing. Concurrent engagement of the parahippocampal gyrus contributes robust contextual and episodic memory integration, while the precentral gyrus provides a streamlined pathway for translating cognitive intentions into precise motor actions. The inclusion of primary auditory cortex further strengthens auditory–linguistic coupling, facilitating rapid transformation of perceptual input into higher-order representations. Collectively, this pattern suggests a functional brain organization that is more tightly coordinated and multimodally integrated than typically observed.

These data suggest that both alleles lead to a humanized brain albeit utilizing different strategies. The direct *CHRFAM7A* allele infers enhanced microstructure consistent with the actin cytoskeleton gain of function ^4^. The inverted *CHRFAM7A* allele leads to higher global efficiency and small world propensity mediated by the microtubule cytoskeleton gain of function.

## Discussion

The human brain has undergone not just quantitative but qualitative evolutionary changes reflected by unique functions. Human specific genes, such as *CHRFAM7A*, can be a contributing force to these qualitative differences. While other human specific genes and genetic variants have been implicated in brain evolution, *CHRFAM7A* is unique as it has two variants with distinct molecular and cellular mechanisms. While the direct allele has been characterized as a modifier of the α7 nAChR, the function of the inverted allele remained elusive.

We engineered isogenic human iPSC to investigate the effect of the inverted *CHRFAM7A* gene carrying the 2 bp deletion in exon 6. We uncovered that inverted *CHRFAM7A* regulated *ULK4* isoform expression via genetic epistasis, resulting in *ULK4* hypermorphism. Long ULK4 is distinctly different from short isoform as it harbors HEAT repeats consisting of repeated pairs of alpha helices that form a solenoid like structure implicated in protein-protein interaction ^14^. As a protein scaffold, ULK4 may play a role in cytoskeletal regulation. ULK4 long isoform is the predominant form in adult brain, with highest expression in the cortex, hippocampus and thalamus ^15^. ULK4 has been implicated in microtubule remodeling, cell motility and neurite branching and elongation ^14^.

Knock-out mouse models and siRNA knockdown of *ULK4* in mice targeted the early exons (shRNA to exons 2-7 and hypomorph mouse model of exon 7) ^17,19^, thus affecting both isoforms and resulting in ULK4 loss of function. In contrast, in the human brain inverted *CHRFAM7A* allele regulates ULK4 through genetic epistasis shifting short to long *ULK4* isoform expression resulting in increased α-tubulin acetylation and peripheral distribution. Thus inverted *CHRFAM7A* allele exerts its effect on ULK4 as an isoform specific gain of function leading to subtle hypermorphism. The skewing from short to long isoform likely expands ULK4’s interaction capacity through the armadillo domain, enabling more complex signaling roles in neurite outgrowth, microtubule stability, and synaptic organization ^36^.

The regulation of *ULK4* isoforms modulates tubulin acetylation in our study; consistent with previous reports describing that depletion of ULK4 in SH-SY5Y cells and primary cortical neurons decreases α-tubulin acetylation ^15^. Tubulin acetylation increases microtubule flexibility and debundling leading to branching ^34,37^. The increased α-tubulin acetylation leads to microtubule cytoskeleton phenotypes (cell body: microtubule rich projections; neurite: debundling; growth cone: fanning with MT invasion). These morphological differences lead to key neuronal functions including neurite outgrowth (primary branching), neurite elongation and branching, phenotypes previously attributed to ULK4 ^15^. ULK4 knockdown in mice demonstrated decreased dendrite length and branching ^38^. In mouse cortex, silencing Ulk4 impairs neuronal arborization, producing abnormal apical dendrites and excessive secondary branching. In knockout mouse brain, dendritic network complexity is reduced ^18^.

In the inverted MGE progenitors we detected an increased number of primary neurites leading to multipolarization in the presence of the inverted *CHRFAM7A* allele (Fig. 4H). In our differentiation protocol the majority of MGE progenitors differentiate into GABAergic neurons ^30^ suggesting a role of inverted *CHRFAM7A* in enhancing GABA interneuron arborization and function. Of note, increased somatic arborization has been described as a strategy to increase computational capacity of the human brain ^14^.

Inverted MGE progenitors demonstrated a dynamic MT cytoskeleton with increased locomotion and GC exploratory movement. These are key mechanisms in neuronal development and plasticity implying a role of the inverted allele in these processes. The increased motility is cell-type independent, detected in iPSC derived and primary human monocytes as well.

Taken together, the *CHRFAM7A* inverted allele associated increased debundling, branching and neurite tip exploration support a hypothesis of a neuronal system with higher connectivity. To test this hypothesis we returned to the human model and performed cognitive, functional and structural imaging on individuals with 0, 1 and 2 copies of the inverted allele. In functional imaging analyses, we found higher node strength, clustering coefficient and small world propensity with increasing copy number of the inverted allele. Increased small-worldness in neural networks is thought to be a functional consequence of the structural properties of neurons, including their arborization and connectivity ^39^. Increased arborization of GABAergic interneurons facilitates local connectivity and network development ^40^, which may account for the observed enhanced small world propensity for the inverted allele, connecting cellular biology to brain organization. Importantly, higher global efficiency and small worldness is associated with cognitive flexibility and adaptability and is implicated in recovery after injury ^41,42^.

In contrast, DTI detected a relative weakness in white matter microstructure as a function of the inverted allele copy number. As the null allele is rare in the human population the 0 copy inverted individuals are homozygous for the direct allele. Thus the difference is likely related to direct allele mediated strength in microstructure. We have shown previously that the direct *CHRFAM7A* allele activates Rac1 small GTPase and leads to a phenotypic switch from filopodia to lamellipodia at every level of the neuron, including cell body, growth cone and dendritic spine^4^. The actin cytoskeleton reinforces the neuronal microstructure and may lead to a more resilient brain. Direct allele associated right-sided enhanced microstructure and inverted allele associated left-predominant strength in functional connectivity may contribute to altered disease susceptibility ^43^. Further larger studies that incorporate measures of brain atrophy, resilience, cognitive reserve and neuronal plasticity in the context of the *CHRFAM7A* alleles are needed to fully understand their impact on the human brain.

The observed binary partitioning of brain organization can be viewed as two distinct cytoskeleton-based strategies that expand neural computational capacity, each driven by a different *CHRFAM7A* allele. The direct *CHRFAM7A* allele promotes an actin-cytoskeleton gain of function, enhancing microstructural refinement through dynamic modulation of dendritic spines and synaptic interfaces. This actin-driven mechanism supports high-resolution local computation, precision encoding, and rapid adaptive plasticity. In contrast, the inverted *CHRFAM7A* allele confers a microtubule-cytoskeleton gain of function, strengthening long-range structural stability, intracellular transport, and large-scale network coordination, thereby enhancing functional connectivity across distributed brain regions. Together, these allele-specific cytoskeletal programs form a binary organizational logic—actin for microcircuit specialization and microtubules for integrative network coherence. Human divergent evolution appears to have converged on the cytoskeleton as a key substrate for boosting neural computation, with the direct and inverted CHRFAM7A alleles representing complementary evolutionary solutions to the challenge of expanding brain function. This binary partitioning of brain organization may infer susceptibility to different diseases and may be conducive to distinct mechanisms. Disease associations reported with the *CHRFAM7A* alleles ^20^ are likely related to the brain organization and disease mechanism interaction. Factoring in this human genetic background may lead to more precise and effective treatments in neuropsychiatric diseases.

## Methods

### Locus specific dual genotyping

Genomic DNA was isolated from whole blood. CHRFAM7A copy number by TaqMan assay: Primers (forward primer: GTAATAG TGTAATACTGTAACTTTAAAATGTGTTACTTGT, reverse primer: AGCCGGGATGGTCTCGAT) and probe (TCCTGACTGTACAC ATAAAA) were designed to detect the breakpoint sequence (Applied Biosystems). The duplex real-time PCR assays were performed using a FAM dye-labeled assay targeted to CHRFAM7A and the VIC dye-labeled RNaseP (TaqMan copy number reference assay, part # 4403326) as a reference gene. Each sample was assayed in quadruplicate by using 10 ng DNA in each reaction. Realtime PCR was performed using the CFX384 Real-time PCR Detection System (Bio-Rad). Threshold cycle (Ct) values were determined for CHRFAM7A and compared with Ct values for RNase P. Relative quantity was determined by the DD Ct method.

CHRFAM7A 2 bp deletion assay: Genotyping for the 2 bp deletion polymorphism was by limited cycle fluorescent PCR (21 cycles of 948C/30 s, 588C/30 s, 728C/1.5 min) using primers flanking the 2 bp deletion. 2bpForFAM GGGCATATTCAAGAGTTCCTGCTAC and 2bpRev. CCACTAGGTCCCATTCTCCATTG gave a product size of 170 bp in the absence of the deletion and 168 bp in its presence. PCR products were resolved using a 3,100 fluorescent genotyper and Genemapper v3.0 software to ascertain 170 bp:168 bp ratios. Each assay was performed at least twice.

### RNA-Seq Gene expression^44^

Bulk RNA sequencing of dorsolateral prefrontal cortex tissue from 205 NCI subjects from the ROSMAP study were utilized for *ULK4* gene expression analysis ^45^. Genotype distribution, age, sex and PMI are depicted in Supplementary Table 1.

Previous reports described the ROSMAP study design and data collection scheme in detail ^46^. Briefly, RNA-seq libraries were generated from high-quality RNA and sequenced on Illumina platforms. Output reads were aligned to the GRCh38 reference genome using STAR in two-pass mode, and abundance of transcripts was quantified using Kallisto with GENCODE annotation. RNA isolation of brain tissue was performed on a subset (48 participants) for ULK4 short and long isoform targeted qPCR assay.

Gene-level for those expressed in more than 50% of samples were used and normalized using conditional quantile normalization (CQN), followed by log2-CPM transformation and voon-based quartile normalization. Lineal regression models including post-morten intervals, sequencing batch, RNA quality metrics, and alignment QC measures were applied to remove technical variability. Normalized expression values were used in downstream analyses.

### Human post mortem brain ULK4 isoform analysis

RNA-Seq raw FASTQ files were splice-aligned to the GRCh38 GENCODE p13 reference genome and r38 annotation using STAR ^47^ (v2.7.9). Multimapping parameters were adjusted appropriately during alignment. The resulting BAM files were then used for isoform quantification with RSEM ^48^ (v1.3.3). Downstream statistical visualizations and analyses were performed using R (https://www.R-project.org) (v4.2.0) statistical software, with plots generated using ggplot2 ^49^ (v3.4.4) package. Isoform quantification, expressed as Transcripts Per Million (TPM) values, was consolidated into a matrix. ULK4 isoforms (ENST00000414606.1, ENST00000420927.5, ENST00000301831.9) were extracted. Violin plots were generated to visualize the distribution of expression for these isoforms, and pairwise scatter plots were created to examine their expression relationships. Samples were matched and labeled with corresponding copy number information. Statistical analysis was performed by Spearman correlation between isoforms.

### Human primary monocytes

#### Source of primary human monocytes

Healthy volunteers meeting inclusion and exclusion criteria signed the informed consent. Inclusion criteria include healthy adult males and females. Exclusion criteria include chronic, autoimmune or systemic inflammatory disease (rheumatoid arthritis, systemic lupus, myositis, inflammatory bowel disease, type 1 diabetes, autoimmune thyroiditis), use of NSAID within 48 hrs of blood draw, immunosuppressant medication use, any concurrent immune based therapies, fever within 36 hrs of blood draw, steroid use for any reason, anticoagulation or antiplatelet therapy, history of bleeding disorder, history of immune deficiency, pregnancy or treatment for cancer, blood donation within the 6 weeks prior to blood draw and use of illegal recreational drugs. Each donor provided up to 60 ml blood which was processed immediately for PBMC isolation. Yield of PBMC was quantified by cell counting and viability recorded. Sample was further processed for monocyte isolation if the viable PBMC was at least 500,000. Due to high human variance of monocyte percentage in humans, the number of monocytes ranged from 20,000-1,750,000. The available amount was prioritized in the following order: DNA isolation/genotyping, RNA isolation/qPCR, ICC and morphological analysis, migration assay.

#### Isolation of primary human monocytes and macrophages differentiation

Peripheral blood was obtained from healthy adults with informed consent, and peripheral blood mononuclear cells were isolated by density gradient centrifugation using SepMate™ PBMC Isolation Tubes (Stem Cell Technologies, Vancouver, Canada) and LymphoPrep (1.077 g ml^−1^; Stem Cell Technologies, Vancouver, Canada) according to the manufacturer’s instructions. Human primary monocytes were further isolated from the peripheral blood mononuclear cells using the Dynabeads® Untouched™ human Monocytes kit from Invitrogen (Carlsbad, CA, USA) according to the manufacturer’s protocol. For further macrophage maturation, monocytes were attached and cultured for 7 days in 10% FBS/RPMI medium supplemented with MCSF (100 ng/ml).

### iPSC model

#### Isogenic lines

Isogenic line development was described previously ^2^. Briefly, we inserted CHRFAM7AΔ2bp coding sequence obtained from Addgene (pcDNA3.1-CHRFAM7AΔ2bp (Addgene plasmid#62515) into the AAVS1 safe harbor site of the constitutively expressed gene PPP1R12C on chr19 (position 19q13.42) of the 0 copy ancestral line (UB068) using TALEN mediated gene editing. The expression vector harbors puromycin resistance gene for selection and fluorescent signal (GFP) for visualization in downstream experiments.

#### Neuronal differentiation

Neuronal differentiation of iPSC towards Medial Ganglionic Progenitors (MGE) was carried out as described previously ^30^. Undifferentiated UB068 (null) and UB068_CHRFAM7A (CHRFAM7AΔ2bp KI) cells grown on irradiated mouse embryonic fibroblasts were detached with Dispase at day 0 and continued to float in T-25 flasks with “iPSC Culture Media” without bFGF. When iPSCs formed Embryoid Bodies (EB) at day 5, the medium was replaced with neural induction medium (NIM) [DMEM/F12+Glutamax, N2 (Life Technologies), NEAA (Gibco), Heparin (Stem Cell Technologies) and pen/strep] on day 4. EBs were attached in 6-well plates on day 7, and by day 10 neural rosettes were present, indicating the development of primitive neuroepithelia. Ventralization of primitive neuroepithelia was started on day 10 by adding Purmorphamine 1.5 µM. On day 16, neural tube-like rosettes were detached and transferred into T-25 Flasks in NIM with 2% B27 (Life Technologies) to form neurospheres. The cultures were fed every other day. On day 25, MGE progenitors were dissociated with Accutase (Stem Cell Technologies) and plated onto 6-well plates for further differentiation in neuronal media.

#### iPSC derived monocyte differentiation

Monocytes/macrophages were differentiated as we described previously according to a well-established protocol ^5,50^. Briefly, monocytes differentiation started with the iPSC detachment (at least two confluent 6-well plates) and embryoid body (EB) formation. At day 4, the EBs were plated in X-Vivo 15 ™ medium (Lonza) supplemented with IL-3 (25 ng/ml) and MCSF (50 ng/ml). From day 15-20, the generated monocytes were harvested twice a week from the supernatant and filtered through a 70 µm mesh nylon strainer (Falcon). For further macrophages maturation, the cells were attached and cultured in X-Vivo 15 ™ medium (Lonza) supplemented with MCSF (100 ng/ml).

### Transfection

Using Ma et al. protocol ^51^ and Lipofectamine 2000 (Thermo Fisher Scientific, Cat# 11668027), UB068 and UB068_Δ2bp MGE progenitors were transfected with 100 nM of Human ULK4 siRNA (Horizon Discovery). Non-targeting Pool siRNA was used as control (Scrambled siRNA). Following transfection, 100,000 cells were plated on 12-mm glass coverslips. For gene expression analysis, the cells were collected 24h after transfection; for ICC – 48h, 72h and 96h after transfection.

### Immunoprecipitation CHRFAM7A

Immunoprecipitation (IP) of the CHRFAM7A and a7 nAChR was performed with Pierce Classic magnetic IP/Co-IP Kit (Thermo Fisher) according to Brai et al ^52^ with modifications. Briefly, MGE progenitors derived from the inverted and the inverted null (UB068_Δ2bp, UB068 and UB068_CHRFAM7A, respectively) lines were plated for 72h. Total cell lysates were obtained using lysis buffer (Thermo Fisher), and protein concentration was determined with Precision Red Advanced Protein Assay (Cytoskeleton Inc). 1 mg of total protein was incubated with 10 μg of anti-CHRFAM7A (Sigma Aldridge, Cat# AV35409) and/or anti-α7nAChR antibodies (Santa Cruz Biotechnology, sc-58607) (overnight, 4°C) and immunoprecipitated using a Protein G Dynabeads (Thermo Fisher). An isotype specific IgG control was used as a negative control. CHRFAM7A and α7 nAChR were detected by Western blot with appropriate antibodies (Sigma Aldridge, Cat#M220).

### Blocking ULK4 antibody

Specificity of ULK4 signal detected by Western blot and by immunocytochemistry, was confirmed by using the specific ULK4 blocking peptide according to manufacturer’s protocol (FabGennix International Inc). Briefly, 10 µl of ULK4 antibody were mixed with 70 µl of blocking peptide or PBS. Following incubation on a rotator (overnight, 4°C) and centrifugation (12,000xg, 2 min, 4°C), both the antibody-peptide and antibody-PBS solutions were diluted to the final volume of 5 ml and used for the downstream application.

### Western blot

Following IP and to detect the amount of ULK4 (FabGennix International, Cat# ULK4-401AP), total (Cell Signaling Technology, Cat#3873T) and acetylated α-tubulin (Sigma Aldridge, Cat# T7451) in the cell lysates collected from MGE progenitors, the samples were separated on 4–20% sodium dodecyl sulfate–polyacrylamide gel electrophoresis (Bio-Rad), transferred onto polyvinylidene difluoride membrane (Bio-Rad), and incubated overnight at 4 °C with primary antibodies (listed in Supplementary Table 12). Next day, the membrane was extensively washed and incubated with appropriate secondary antibodies. Specific immunoreactive bands were detected using ChemiDoc XRS + Imaging Systems (Bio-Rad). Densitometry analysis was performed using Image Lab 6.0.1 Software. GAPDH was used as a loading control.

### Immunocytochemistry and confocal microscopy

MGE progenitors and/or neurons plated on Matrigel-coated glass coverslips or 8-well glass chambers (Thermo Fisher) were fixed with 4% paraformaldehyde (Mallinckrodt Baker) for 15 minutes, permeabilized with 0.1% Triton X100 (Mallinckrodt Baker) for 10 minutes and blocked with blocking buffer (5% BSA in PBS) for 1 hour at room temperature (RT). Cells were incubated overnight at 4°C with primary antibodies and for 1h at RT with secondary antibodies. Both primary and secondary antibodies were diluted in a blocking buffer. Actin was visualized with ActinRed 555 ReadyProbes (Thermo Fisher Scientific). Slides/coverslips were incubated with NucBlue (DAPI) for 5 min at RT and mounted with a ProLong® Gold Antifade reagent (Life Technologies). Confocal images were captured by using Leica TCS SP8 confocal microscope system (63x and/or 20x objective). Images were acquired using LAS X software.

### Image Analysis

#### ULK4 distribution quantification

Cellular localization of ULK4 (the nuclear/cytoplasmic ratio) was quantified as described previously ^31^. Following ICC, the confocal images were taken a 63x objective, and the mean florescent signal of ULK4 in the nucleus (DAPI positive area) and in the cytoplasm was quantified using Fiji ImageJ in at least 10 images/sample (cell line) in the three independent experiment.

### RNA isolation from human brain

RNA was isolated from human brain samples using Lysing Matrix D (MP Biomedicals). Briefly, 20-30 µg of brain tissue immersed in Trizol (Invitrogen) was transferred to Lysing Matrix D tube and vortexed 5-7 times till the beads mostly dissolved. Afterwards, RNA was isolate according to Trizol isolatiol protocol.

### RT PCR

Total RNA was isolated using Trizol (Invitrogen) according to manufacturer’s protocol. cDNA was synthesized from 1µg RNA using ImProm-II reverse transcriptase (Promega) and oligo (DT) (Promega) at 42°C for one hour.

### Long and short ULK4 specific amplification

PCR amplification of template of the short ULK4 isoform was performed using specific exon11 forward (5’-CAACTGAGTTTCGGCCTAAGAG-’3) and exon 18short reverse (5’-ACAAGGCACTGTCTAGGCAC-’3) primer pair, for the long isoform amplification of template was performed using specific exon11 forward (5’-CAACTGAGTTTCGGCCTAAGAG-’3) and exon19 reverse (5’-CGAAGGCACCTCATTAGCAC-3’).

### Isoform specific sequencing

cDNA corresponding to ULK4 short and long isoforms was amplified using a exon 11 forward primer paired with exon 18 reverse primer specific for short isoform and exon 19 reverse primer for long isoform. The resulting PCR products were sequenced by Sanger sequencing, and sequence alignments were conducted using Clustal O.

### Predicted secondary structure of mRNA

The predicted secondary structure of short and long mRNA structures were obtained from GENECODE/UCSC database (https://genome.ucsc.edu/).

### qPCR

Quantitative gene expression was detected by standard RT-qPCR using specific primers (IDT) listed in Supplementary Table 11 and SYBR green master mix (Biotool) run on Bio-Rad CFX Connect cycler (Bio-Rad). Relative gene expression was quantified by ΔΔCT method normalized to GAPDH expression assayed with 3 technical replicates.

### Migration assay

Migration of the iPSC-derived and PBMC-derived monocytes was assessed using CHEMICON® QCM™ 24-well Migration Assay Kit (MilliporeSigma) according to the manufacturer’s instructions. 200,000 cells were plated in the upper chamber of the insert with an 8μm pore size and allowed to migrate for 16h. CXCL12 (100 ng/ml) was used as a chemoattractant. Migration was detected colorimetrically measuring absorbance at 560 nm.

### Live imaging

MGE progenitors generated from UB068 and UB068_Δ2bp cell lines and human primary monocytes were plated on 8-well chambers (30, 000 cells/well) and transfected with 0.5 μg of Lifeact-RFP according to Ma et al. protocol ^51^ for live actin imaging or with SPY555-tubulin dye for 1 hour (SpiroChromes, CY-SC203). Live imaging was performed 16h after transfection or 1 hour after dye staining using Leica DMi8 microscope (objective 63x). The cells were kept at 5% CO2 and 37°C. The images were acquired every 4 sec for 5 min. At least 10 cells/ line/experiment were imaged in four independent experiments.

### Image analysis and quantification

Images acquired were analyzed in imageJ.

#### Neurite length

NeuronJ plugin was used for semiautomatic tracing of neurites in MGE progenitors stained for acetylated α-tubulin. Mean length was quantified and the length of each neurite was summed to obtain the total length per cell.

#### Sholl analysis

MGE progenitors stained for acetylated α-tubulin were individualized by creating a binary image to visualize the cell skeleton. Sholl Analysis of Neuroanatomy plugin was performed creating concentric circles in the center of the cells with a step size of 9 µm and a end radius of 215 µm to measure total intersections.

#### Neurite classification

processes were classified in axons/primary neurites, secondary neurites and tertiary and each type of process was compared between cell lines.

#### Neutite tracking

regions of interest (ROIs) sized 10 µm x 10 µm were selected. Binary images were obtained and MTrackJ plugin was used to select the neurite tip overtime and measure the total distance traveled and distance from the starting point. For <u>kymographs</u>: kymographs were created drawing a 13 µm segments perpendicularly to the microtube bundles to create a 2D projection of the time course.

#### Neurite microtubule debundling

segments were used to measure the max diameter of bundles in axons of MGE progenitors stained with SPY555-tubulin. On average, 9-10 images were taken per experiment in four independent experiments.

### Participants in cognitive and imaging study (CEG-MS study)

University at Buffalo Institutional Review Board (IRB) approved the study, study ID 030–603069. Informed consent procedure was performed. The healthy participants in this sub-analysis were part of a larger prospective study that aimed at investigating the cardiovascular, environmental and genetic risk factors in multiple sclerosis (CEG-MS study) ^53^. Inclusion criteria included (1) age 18–75, (2) willingness to participate in neuropsychological, MRI and clinical investigation. Exclusion criteria included (1) history of any major neurological or psychiatric diagnosis, (2) use of psychoactive medications, (3) pregnant or nursing mothers, and (4) any contraindications in completing the study procedures (i.e., MRI examination). All participants provided signed consent form.

### Motor and cognitive battery (CEG-MS study)

Subjects underwent neuropsychological testing which included the oral response version of the Symbol Digit Modalities Test (SDMT),^54,55^ measuring cognitive processing speed (also visual scanning working memory/learning and lexical access speed), and targeted tests for auditory/verbal learning and memory (California Verbal Learning Test – 2nd Edition or CVLT+II^56^), and visual/spatial learning and memory (Brief Visuospatial Memory Test – Revised; BVMT-R^57,58^). The battery was administered by a trained technicians under supervision of board-certified neuropsychologist (RHBB). In all aforementioned cognitive measures, higher score indicates better cognitive performance. Depression and fatigue were quantified by the Beck Depression Inventory – Fast Screen (BDI-FS)^59^ and the Fatigue Severity Scale (FSS)^60^ to exclude confounding effect. In both patient-reported outcomes, higher scores indicate worse symptoms.

### Personality assessment (CEG-MS and BLSA studies)

The CEG-MS study assessed Big Five personality traits using the NEO Five-Factor Inventory (NEO-FFI), a 60-item self-report questionnaire that provides brief domain-level scores for Neuroticism, Extraversion, Openness, Agreeableness, and Conscientiousness. The Baltimore Longitudinal Study of Aging (BLSA) used the Revised NEO Personality Inventory (NEO-PI-R), a 240-item instrument that offers a more detailed assessment of the same five personality domains, including facet-level subscales. The T scores for Neuroticism, Extraversion, Openness, Agreeableness, and Conscientiousness were normalized to population means of each instrument. The normalized scores were combined for statistical analysis.

### MRI acquisition (CEG-MS study)

All healthy participants were scanned using the same 3 T GE Signa Excite Scanner (GE, Milwaukee, WI) and 8-channel head and neck coil. The sequences that are relevant for the analysis in this study were: (1) 3D high-resolution T1-weighted inversion recovery fast spoiled gradient echo with echo time (TE) of 2.8 ms, repetition time (TR) of 5.9 ms and inversion time of 900 ms, flip angle of 10 degrees, field-of-view of 25.6 × 19.2 cm2 and isotropic 1 × 1 × 1 mm slices, (2) functional MRI (fMRI) that acquired 240 volume of gradient echo-echo planar images with TE of 35 ms, TR of 2,500 ms, flip angle of 90 degrees and 3.75 × 3.75 × 4 mm slices and (3) fluid-attenuated inversion recovery (FLAIR) sequence with TE of 120 ms, TR of 8,500 and TI of 2,100 ms, flip angle of 90 degrees, echo train length 1 × 1 × 3 mm slices with no gap. There were no software changes during the acquisition of all participants.

### MRI processing

The T2 lesion volume (LV) was determined on FLAIR scans by experienced neuroimager using semi-automated contouring and thresholding tool and corrected with the Java Image Manipulation software (JIM, Xinapse systems, Essex, UK, version 8.0) ^61^. All T1 weighted images were preprocessed for N4 bias field correction and lesion inpainting. The segmentation of the brain volumes including the whole brain volume (WBV), white matter volume (WMV), gray matter volume (GMV), lateral ventricular volume (LVV), deep gray matter volume (DGMV) and thalamic volume were performed using the cross-sectional Structural Image Evaluation, using Normalization, of Atrophy (SIENAX1) ^62^ and FMRIB’s Integrated Registration and Segmentation Tool (FIRST2) ^63^ protocols. All volumes were normalized for the head size. The cortical parcellation was performed using the FreeSurfer protocol that provides the cortical map of 86 regions based on the Desikan-Killiany atlas3 ^64^.

#### Diffusion-weighted imaging preprocessing

The topup tool, part of the FMRIB’s Software Library (FSL) toolbox (http://www.fmrib.ox.ac.uk/fsl), was used to correct for susceptibility-based geometric ^65^. As reversed phase-encoding b=0 images were not available, we used the Synb0-DisCo method to generate a synthetic, distortion-free b=0 image from the original b=0 image along with the 3D T1-weighted image ^66^. Eddy current distortions and subject movement were corrected by using the eddy tool ^67^, which is also part of the FSL toolbox. The diffusion tensor was then fitted, and fractional anisotropy (FA), mean diffusivity (MD), axial diffusivity (AD), and radial diffusivity (RD) maps were obtained.

#### White matter analysis

The tract-based spatial statistics (TBSS) pipeline ^68^ was then utilized to generate skeletonized WM maps. All subjects’ FA data were aligned into a common space using nonlinear registration. WMLs were excluded from the registration cost function to minimize their impact on spatial normalization. Next, the mean FA image was created and thinned to create a mean FA skeleton which represents the centers of all tracts common to the group. Each subject’s aligned FA, AD, MD, and RD data were then projected onto this skeleton.

Joint inference on FA, AD, MD, and RD maps was performed using the non-parametric combination (NPC) method ^69^ as implemented in the Permutation Analysis of Linear Models (PALM) tool ^70^. With NPC, the joint null hypothesis of the NPC is that the null hypothesis for each of the partial tests (i.e. univariate analysis of the individual modalities) is true while the alternative is that any of them are false. It is important to note that the NPC technique does not simply take the union of any suprathreshold voxels but instead combines the partial tests into a joint statistic. Consequently, NPC offers a more powerful alternative compared to using just a single modality or performing post-hoc analyses of multiple modalities. General linear models (GLM) were tested with PALM using tail acceleration ^71^ with 500 permutations and threshold-free cluster enhancement (TFCE) for computing spatial statistics ^72^, adjusting for age, sex, and diffusion acquisition protocol. For the NPC analyses, partial tests were combined using the default Fisher function.

### Resting State functional Magnetic resonance imaging (CEG-MS study)

The resting-state fMRI was processed using FSL tools as described elsewhere, following Human Connectome Project preprocessing recommendations ^73^. Briefly, the processing included removal of the first 2 volumes, slice timing correction, motion correction, intensity normalization, high-pass temporal filtering (2,000 s), field map unwarping based on phase-reversed acquisitions (blipup/down), and 4-mm spatial smoothing. Motion confounds (of the 6 rigid-body parameter timeseries), cerebrospinal fluid signal, and white matter signal were regressed out. The activity between regions and their functional connectivity was determined by assessing the concordance of temporal activation using partial correlation estimation using Nilearn ^74^. Matrices of the 86 × 86 regions and their paired functional connectivity were produced.

### Graph theory analysis

Functional connectivity measures of the parcellated were derived from the aforementioned thresholded 86 x 86 functional connectivity matrices using the Brain Connectivity Toolbox (BCT4) ^75^ and custom MATLAB software ^76^. The functional connectivity values were Z-transformed prior to graph analysis. Positive- and negative-correlation networks were analyzed separately. Selected network measures included average node strength, global efficiency, average global clustering coefficient, average global path length, and small world propensity.

### Statistical analysis

All experiments were performed in triplicates or quadruplicates and values are expressed as mean ±SD. Averages were compared to determine significant differences by unpaired parametric two sided t-test, unpaired non-parametric with Mann-Whitney U-test and one-way ANOVA for multiple comparisons followed by Tukey test. Comparisons between cumulative density functions were applied by the nonparametric two-sample Kolmogorov–Smirnov test. For Spearman rank correlation averages were compared by two-tailed t-test with Bonferroni as post-hoc test. All statistical analyses were performed using R software package (version 4.1.2) and GraphPad Prism (version 10.1.0).

Power calculation was not performed due to the exploratory nature of this clinical-imaging-genetic correlation study thus sample size was not predetermined. Significance was set at p < 0.05. Additional correction for false discovery rate (FDR) using the Benjamini-Hochberg procedure was performed and FDR-corrected p-values were also shown. The data distribution was determined by visual inspection of the histograms and Q-Q plots. Age was compared using analysis of variance (ANOVA). The non-parametric data was compared using Mann Whitney U test. Comparison between CHRFAM7A carriers and non-carriers for cognitive data were age and years of education-adjusted using analysis of covariance (ANCOVA) and depicted as estimated marginal means (standard error). MRI measures were compared by age-adjusted analysis of covariance (ANCOVA) and described as mean (standard error) and as estimated marginal means corrected for age. SPSS (Armonk, NY, United States) version 28 statistical software was used for all analyses and GraphPad Prism (San Diego, CA, United States) was used for data visualization and creation of the data plots.

## Supplementary information

Supplementary Figures 1-2

Supplementary Videos 1-2 Supplementary Tables

## Limitations

Cell culture has its limitations as it cannot model the temporal-spacial-functional complexities of the brain. However, isogenic human iPSC provides the human context that we cannot achieve with model organisms. An inverted *CHRFAM7A* KI mouse model or human iPSC organoids can help further characterize the phenotypes at the organ level. Of note, the human cognitive-structural-functional readout is consistent with extrapolation of the iPSC findings, correlating increased small worldness with increased arborization specifically of GABA interneurons. Larger clinical studies on cognitive function, structural and functional neuroimaging, neurophysiology and measures of brain resilence and plasticity in the context of the biallelic human genetic background will likely yield relevant insights.

## Resource availability

Further requests for resources, reagents, or data should be directed to and will be fulfilled by Kinga Szigeti.

All unique/stable material generated in this study are available from the lead contact with a completed materials transfer agreement.

## Acknowledgements

This work is supported in part by the Community Foundation for Greater Buffalo (Kinga Szigeti). ROSMAP is supported by NIA grants P30AG10161, P30AG72975, R01AG15819, R01AG17917. U01AG46152, and U01AG61356. We thank Luigi Ferrucci, Susan Resnick and Juan Troncoso and the Baltimore Longitudinal Study of Aging (BLSA) for providing the postmortem brain samples and personality clinical data used in this study and providing feedback on the manuscript.

