## Supplemental Information for "Binary partitioning of human brain organization due to divergent human cytoskeletal evolution"

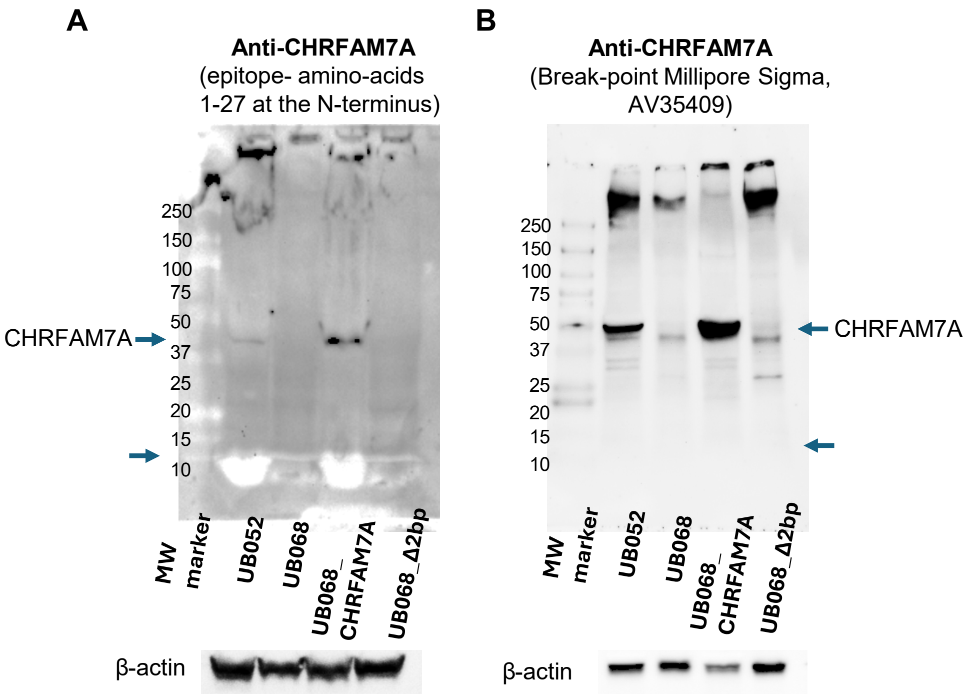


**Supplementary Figure 1.** Detection of CHRFAM7A in iPSC-derived MGE progenitors. A. Membrane probed with the anti-CHRFAM7A antibody, a generous gift from Dr. A. Baird (UCSD) - final bleed, hence used in high dilutions and shows membrane impurities (left panel). The anti- CHRFAM7A antibody (from A. Baird) detects a unique 27 amino acid sequence at the N-terminus, which is not present in CHRNA7. B. Membrane probed with anti-CHRFAM7A antibody (Break-point antibody, Millipore Sigma, AV35409); epitope - first 100aa of the fusion FAM7A and CHRNA7 (right panel). β-actin is used as loading control. Note that neither antibody detects the band in the inverted UB068_Δ2bp line (46kDa or 12kDa). UB052 is a nascent CHRFAM7A direct carrier line.

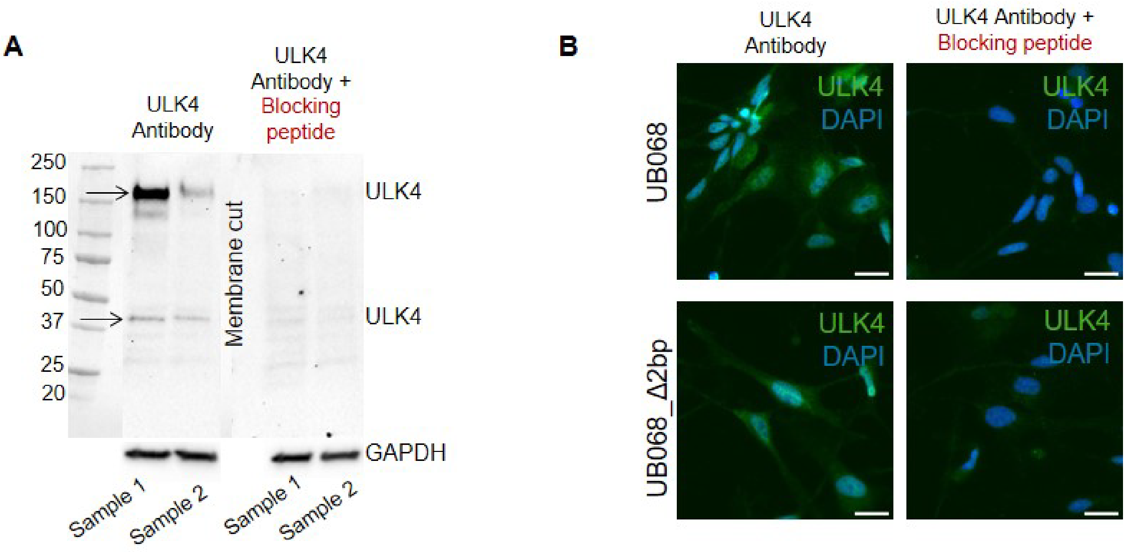


**Supplementary Figure 2. A.** Detection of ULK4 in human macrophages by Western blot Note the presence of two specific bands at 150 kDa and 40 kDa (left part of the blot) that disappear after incubation with ULK4-specific blocking peptide (right part of the blot). **B.** Immunofluorescent detection of ULK4 in the null and inverted MGE progenitors with and without blocking peptide. Scale bar, 20 µm.

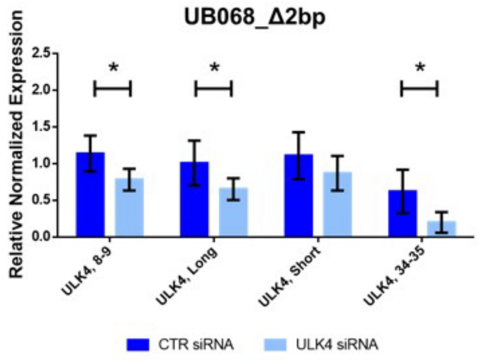


**Supplementary Figure 3.** In MGE progenitors derived from the inverted line, transfection with ULK4 siRNA decreases expression of early and late exons of *ULK4* as well as the Long isoform of *ULK4*. Expression of the Short isoform remains unchanged. Data are means ±SD; UB068_Δ2bp ULK4 Exon 8-9, ULK4 Long, ULK4 Exon 34-35 * $p < 0.05$ multiple t-test using Holm-Sidak method.

***Supplementary video 1:*** Representative videos of human primary monocytes MA033 and MA031stained with SPY555-tubulin during a 304 sec time lapse corresponding to 76 images.

***Supplementary video 2:*** Representative videos of neurites of UB068 and UB068_Δ2bp stained with SPY555-tubulin showing tubulin bundles in a 304 sec time lapse corresponding to 76 images. ROIs are 20x10 μm.

**Supp. Table 1.** Autopsy Cohort Demographics (ROSMAP).

| | **Whole Cohort (n = 205)** | **Non-Carriers (n = 59)** | **1 Inverted Allele (n = 100)** | **2 Inverted Alleles (n = 46)** | **F-value; p-value** |
|---|---|---|---|---|---|
| Age (years)*, mean ± SD | 84.1 ± 5.5 | 83.7 ± 5.5 | 84.0 ± 5.6 | 84.9 ± 5.3 | F = 2.398; p = 0.094[a] |
| Sex* | 79 male, 122 female | 19 male, 38 female | 42 male, 57 female | 18 male, 27 female | p = 0.666 [b] |
| PMI*, mean ± SD | 7.7 ± 4.7 | 7.2 ± 4.2 | 7.4 ± 4.3 | 9.0 ± 5.7 | F = 0.562; p = 0.571[a] |

*4 subjects had missing demographic information.
Abbreviations: SD = standard deviation.
[a]ANOVA.
[b]Chi-squared test.

**Supp. Table 2.** Primary Monocyte Donor Cohort Demographics (UB).

| | **Whole Cohort (n = 43)** | **Non-Carriers (n = 14)** | **1 Inverted Allele (n = 15)** | **2 Inverted Alleles (n = 14)** | **F-value; p-value** |
|---|---|---|---|---|---|
| Age (years), mean ± SD | 39.1 ± 15.8 | 38.6 ± 17.5 | 34.8 ± 9.4 | 44.2 ± 18.9 | F = 1.311; p = 0.281[a] |

| | | | | | |
|---|---|---|---|---|---|
| Sex | 19 male, 24 female | 8 male, 6 female | 5 male, 10 female | 6 male, 8 female | p = 0.432 [b] |

Abbreviations: SD = standard deviation.
[a]ANOVA.
[b]Chi-squared test.

**Supp. Table 3.** CEG-MS Cohort Demographics.

| | **Whole Cohort (n = 46)** | **Non-Carriers (n = 15)** | **1 Inverted Allele (n = 22)** | **2 Inverted Alleles (n = 9)** | **F-value; p-value** |
|---|---|---|---|---|---|
| Age at Baseline (years) , mean ± SD | 45.8 ± 14.9 | 49.2 ± 13.5 | 46.3 ± 12.6 | 54. 8 ± 22.3 | F = 0.487; p = 0.618 [a] |
| Time to follow-up (years) , mean ± SD | 5.5 ± 0.5 | 5.7 ± 0.6 | 5.5 ± 0.4 | 5.4 ± 0.6 | F = 0.821; p = 0.447 [a] |
| Sex | 32 female, 14 male | 9 female, 6 male | 17 female, 5 male | 6 female, 3 male | p = 0.522[b] |
| Education (years) , mean ± SD | 14.6 ± 2.4 | 14.9 ± 2.9 | 13.9 ± 1.8 | 16.0 ± 2.6 | F = 2.609; p = 0.085 [a] |

Abbreviations: SD = standard deviation.
[a]ANOVA.
[b]Chi-squared test.

**Supp. Table 4.** Baltimore Longitudinal Study of Aging Cohort Demographics.

| | **Whole Cohort** | **Non-Carriers** | **1 Inverted Allele** | **2 Inverted Alleles** | **F-value; p-value** |
|---|---|---|---|---|---|
| Number of visits, median [range] | 3 [1, 7] | 2 [1, 5] | 2 [1, 7] | 4 [1, 5] | p = 0.154[c] |
| NEO PI-R | N = 53 | N = 20 | N = 20 | N = 13 | |

| | | | | | |
|---|---|---|---|---|---|
| Age at Baseline (years), mean ± SD | 72.7 ± 11.6 | 71.2 ± 13.1 | 74.3 ± 9.9 | 72.4 ± 12.3 | F = 0.341; p = 0.712[a] |
| Sex | 20 female, 33 male | 7 female, 13 male | 8 female, 12 male | 5 female, 8 male | p = 0.817[b] |
| Number of visits, median [range] | 4 [1, 12] | 3.5 [1, 6] | 3 [1, 10] | 4 [1, 12] | p = 0.227[c] |

Abbreviations: NEO PI-R = NEO Personality Index- Revised; SD = standard deviation.
[a]ANOVA.
[b]Chi-square test.
[c]Independent-samples Kruskal-Wallis Test.

**Supp. Table 5.** Comparison of cognitive testing performance between inverted genotypes (CEG-MS Cohort).

| | 1 copy vs noncarrier[a] | 2 copies vs noncarrier[a] |
|---|---|---|
| BVMT-R Total | B = 2.090 [-1.838 6.017]; p = 0.297 | **B = -5.578 [-10.307, 0.850]; p = 0.021** |
| BVMT-R DR | B = 0.118 [-1.522, 1.758]; p = 0.887 | B = -1.576 [-3.551, 0.398]; p = 0.118 |
| SDMT | B = -0.529 [-7.584, 6.525]; p = 0.883 | B = -3.313 [-11.806, 5.181]; p = 0.445 |
| FFS | B = 0.216 [-0.630, 1.062]; p = 0.616 | B = 0.556 [-0.463, 1.574]; p = 0.285 |
| BDI-FS | B = -0.060 [-1.021, 0.900]; p = 0.902 | B = -0.859 [-2.015, 0.298]; p = 0.146 |
| 9HPT | B = -0.973 [-4.554, 2.607]; p = 0.594 | B = -0.320 [-4.631, 3.990]; p = 0.884 |

N: 15 non-carriers, 20 single inverted copy carriers, 9 two inverted copy carriers.
Abbreviations: 9HPT = 9-hole peg test; BDI = Beck's Depression Inventory- Fast Screen; BVMT-R = Brief Visuospatial Memory Test- Revised; FFS = Fatigue Severity Scale; SDMT = Symbol Digit Modalities Test.
[a]Linear models correcting for baseline age, sex, and brain volume.

**Supp. Table 6.** Comparison of Big Five personality traits between inverted genotypes (CEG-MS Cohort and BLSA).

| **Trait** | Genotype by study interaction[a] | 1 inverted allele vs noncarrier[b] | 2 inverted alleles vs noncarrier[b] |
|---|---|---|---|
| Neuroticism | F = 0.063; p = 0.939 | B = -1.582 [-4.298, 1.134]; p = 0.253 | B = -1.172 [-4.139, 1.050]; p = 0.242 |
| Extraversion | F = 0.088; p = 0.915 | B = 2.961 [0.255, 5.668]; p = 0.032 | B = 2.415 [-0.419, 5.249]; p = 0.095 |

| | | | |
|---|---|---|---|
| Openness | F = 0.028; p = 0.972 | **B = 3.182 [0.353, 6.012]; p = 0.028** | **B = 7.153 [4.191, 10.116]; p < 0.001** |
| Agreeableness | F = 0.389; p = 0.678 | B = 2.226 [ -0.210, 4.661]; p = 0.073 | B = 0.119 [-2.431, 2.668]; p = 0.927 |
| Conscientiousness | F = 0.333; p = 0.717 | B = -0.455 [-3.263, 2.354]; p = 0.750 | B = 2.758 [-0.182, 5.699]; p = 0.066 |

N: 34 non-carriers (88 visits), 41 single inverted copy carriers (107 visits), 22 two inverted copy carriers (87 visits).
[a]Mixed-effects linear models with covariates of study (CEG or BLSA), study by genotype interaction, baseline age, and sex, and random effect of subject.
[b]Mixed-effects linear models with covariates of study (CEG or BLSA), baseline age, and sex, and random effect of subject.

**Supp. Table 7.** Comparisons of baseline brain volumes between inverted genotypes (CEG-MS Cohort).

| | 1 inverted copy vs Non-carrier | 2 inverted copies vs Non-carrier |
|---|---|---|
| Normalized BPV (mL) | B = -2.8E4 [-6.8E4, 1.1E4]; p = 0.157 | B = -2.1E4 [-7.0E4, 2.9E4]; p = 0.412 |
| Normalized GM (mL) | B = -8.8E3 [-3.2E4, 1.5E4]; p = 0.463 | B = 5.8E3 [-2.4E4, 3.5E4]; p = 0.702 |
| Normalized WM (mL) | B = -2.0E4 [-4.4E4, 4.9E3]; p = 0.116 | B = -2.6E4 [-5.7E4, 4.2E3]; p = 0.091 |
| Normalized Cortical GM (mL) | B = -5.0E3 [-2.6E4, 1.6E4]; p = 0.645 | B = 8.6E3 [-1.8E4, 3.5E4]; p = 0.519 |
| Normalized Ventricular Volume (mL) | B = -7.2E2 [-9.5E3, 8.0E3]; p = 0.872 | B = -3.1E3 [-1.4E4, 7.7E3]; p = 0.580 |

*N: 15 non-carriers, 21 single inverted copy carriers, 9 two inverted copy carriers.
Abbreviations: BPV = brain parenchymal volume; GM = gray matter; WM = white matter.
[a]Linear models correcting for baseline age and sex.

**Supp. Table 8.** Tract-Based Spatial Statistics with significant clusters of p < 0.05 (CEG-MS Cohort).

| **Cluster number** | **Cluster extent (number of voxels)** | **Peak MNI coordinate** | **Anatomical region** |
|---|---|---|---|
| 1 | 3667 | 26, -55, 27 | Right inferior fronto-occipital fasciculus |

| | | | |
|---|---|---|---|
| 2 | 393 | 31, -14, 24 | Right superior longitudinal fasciculus |
| 3 | 27 | 20, -15, 42 | Right corticospinal tract |

**Supp. Table 9.** Comparison of graph theoretical network measures between inverted genotypes (CEG-MS Cohort).

| | **Positive-correlation Network[a]** | **Negative-Correlation Network[a]** |
|---|---|---|
| **Average Node Strength** | 1-copy vs noncarrier: B = 0.000 [-0.032, 0.032]; p = 0.986<br><br>**2-copies vs noncarrier: B = 0.057 [0.006, 0.108]; p= 0.030** | 1-copy vs noncarrier: B = -0.001 [-0.029, 0.028]; p = 0.972<br><br>2-copies vs noncarrier: B = 0.035 [-0.008, 0.079]; p= 0.109 |
| **Global Efficiency** | 1-copy vs noncarrier: B = 0.002 [-0.007, 0.012]; p = 0.611<br><br>2-copies vs noncarrier: B = 0.015 [-0.001, 0.031]; p = 0.071 | 1-copy vs noncarrier: B = -0.002 [-0.008, 0.012]; p = 0.746<br><br>**2-copies vs noncarrier: B = 0.020 [0.002, 0.038]; p = 0.029** |
| **Average Clustering Coefficient** | 1-copy vs noncarrier: B = -0.007 [-0.032, 0.018]; p = 0.592<br><br>**2-copies vs noncarrier: B = 0.060 [0.016, 0.105]; p = 0.008** | 1-copy vs noncarrier: B = -0.011 [-0.034, 0.012]; p = 0.332<br><br>**2-copies vs noncarrier: B = 0.042 [0.003, 0.080]; p = 0.034** |
| **Average Path Length** | 1-copy vs noncarrier: B = -0.113 [-0.401 0.176]; p = 0.444<br><br>2-copies vs noncarrier: B = -0.435 [-0.896, 0.025]; p = 0.064 | 1-copy vs noncarrier: B = -0.121 [-0.404, 0.162]; p = 0.401<br><br>2-copies vs noncarrier: B = -0.402 [-0.876, 0.072]; p = 0.097 |
| **Small World Propensity** | 1-copy vs noncarrier: B = -0.016 [-0.034, 0.002]; p = 0.084<br><br>**2-copies vs noncarrier: B = 0.024 [0.003, 0.045]; p = 0.023** | **1-copy vs noncarrier: B = -0.022 [-0.038, -0.006]; p = 0.006**<br><br>2-copies vs noncarrier: B = -0.013 [-0.033, 0.007]; p = 0.209 |

N: 13 non-carriers, 18 single inverted copy carriers, 6 two-inverted copy carriers.
[a]Linear models correcting for baseline age and sex.

**Supp. Table 10.** Brain regions with positive and negative node strength (CEG-MS Cohort).

| | Positive Network | Negative Network |
|---|---|---|

| | | |
|---|---|---|
| Left Middle Temporal Cortex | 1 copy vs noncarrier:<br>B = 0.547 [-1.112, 2.205]; p = 0.518<br><br>2 copies vs noncarrier: B = 4.108 [1.873, 6.343]; p < 0.001 | -- |
| Left Parahippocampal Gyrus | 1 copy vs noncarrier:<br>B = 1.352 [-0.959, 3.664]; p = 0.252<br><br>2 copies vs noncarrier: B = 5.024 [1.909, 8.139]; p = 0.002 | -- |
| Left Precentral Gyrus | 1 copy vs noncarrier:<br>B = 0.643 [-0.900, 2.185]; p = 0.414<br><br>2 copies vs noncarrier: B = 2.152 [0.073, 4.231]; p = 0.042 | 1 copy vs noncarrier:<br>B = 0.564 [-0.181, 1.310]; p = 0.138<br><br>2 copies vs noncarrier: B = 1.451 [0.446, 2.455]; p = 0.005 |
| Left Caudal Anterior Cingulate Cortex | 1 copy vs noncarrier:<br>B = 1.532 [-0.680, 3.744]; p = 0.175<br><br>2 copies vs noncarrier: B = 4.608 [1.627, 7.589]; p = 0.002 | -- |
| Right Caudal Anterior Cingulate Cortex | 1 copy vs noncarrier:<br>B = 1.751 [-0.121, 3.623]; p = 0.067<br><br>2 copies vs noncarrier: B = 2.691 [0.168, 5.213]; p = 0.037 | -- |
| Right Transverse Temporal Gyrus | 1 copy vs noncarrier:<br>B = 0.301 [-1.065, 1.667]; p = 0.666<br><br>2 copies vs noncarrier: B = 2.161 [0.320, 4.001]; p = 0.021 | -- |
| Left Caudate | 1 copy vs noncarrier:<br>B = 0.212 [-1.825, 2.249]; p = 0.838 | -- |

| | | |
|---|---|---|
| | 2 copies vs noncarrier: B = 3.526 [0.781, 6.271]; p = 0.012 | |
| Left Posterior Cingulate Cortex | -- | 1 copy vs noncarrier:<br>B = 0.428 [-1.932, 2.789]; p = 0.722<br><br>2 copies vs noncarrier: B = 3.827 [0.646, 7.008]; p = 0.018 |
| Left Superior Parietal Cortex | -- | 1 copy vs noncarrier:<br>B = 1.449 [-1.465, 4.364]; p = 0.330<br><br>2 copies vs noncarrier: B = 4.731 [0.804, 8.659]; p = 0.018 |
| Left Supramarginal Gyrus | -- | 1 copy vs noncarrier:<br>B = -1.188 [-3.471, 1.095]; p = 0.308<br><br>2 copies vs noncarrier: B = 4.346 [1.269, 7.423]; p = 0.006 |
| Right Cuneus | -- | 1 copy vs noncarrier:<br>B = -0.070 [-2.573, 2.432]; p = 0.956<br><br>2 copies vs noncarrier: B = 5.467 [2.095, 8.839]; p = 0.001 |
| Right Inferior Temporal Cortex | -- | 1 copy vs noncarrier:<br>B = -0.328 [-2.038, 1.383]; p = 0.707<br><br>2 copies vs noncarrier: B = -2.703 [-5.009, -0.398]; p = 0.022 |
| Right Supramarginal Gyrus | -- | 1 copy vs noncarrier:<br>B = -0.427 [-2.724, 1.871]; p = 0.716<br><br>2 copies vs noncarrier: B = 4.707 [1.611, 7.804]; p = 0.003 |
| Right Insula | -- | 1 copy vs noncarrier: |

| | | |
|---|---|---|
| | | B = 0.887 [-0.827, 2.600]; p = 0.310<br><br>2 copies vs noncarrier: B = 2.375 [0.067, 4.684]; p = 0.044 |
| Left Pallidum | -- | 1 copy vs noncarrier:<br>B = -0.737 [-3.211, 1.737]; p = 0.559<br><br>2 copies vs noncarrier: B = 4.262 [0.928, 7.596]; p = 0.012 |
| Right Thalamus | -- | 1 copy vs noncarrier:<br>B = -0.507 [-2.326, 1.312]; p = 0.585<br><br>2 copies vs noncarrier: B = -3.855 [-6.306, -1.404]; p = 0.002 |
| Right Caudate | -- | 1 copy vs noncarrier:<br>B = -1.054 [-2.721, 0.613]; p = 0.215<br><br>2 copies vs noncarrier: B = -2.803 [-5.049, -0.557]; p = 0.014 |

**Supp. Table 11.** Primer Sequences used for RT-qPCR

| Gene | Forward Sequence | Reverse Sequence |
|---|---|---|
| GAPDH<br>ULK4 - long<br>(Ex.17-18Long)<br>ULK4 – short<br>(Ex.17-18Short)<br>ULK4 (Ex 8-9)<br>ULK4 (Ex 34-35)<br>CHRNA7[1]<br>CHRFAM7A[1] | GTTCGACAGTCAGCCGCATC<br>GCCAAGGTTGCTCACGTAAT<br>GCCAAGGTTGCTCACGTAAT<br>CTTCTCGTCCTA<br>AAGCTTCTTCA<br>TGATTAGCCTGCTCATTCCA<br>ACATGCGCTGCTCGCCGGGA<br>ATAGCTGCAAACTGCGATA | GGAATTTGCCATGGGTGGA<br>AGGGTTGGTAAAAGGCACTG<br>CACAAGGCACTGTCTAGGCA<br>GATCTTCGACGCTTGATTCC<br>CTTCTGCTCCTTTGGGTC CT<br>GATTGTAGTTCTTGACCAGCT<br>CAGCGTACATCGATGTAGCAG |

[1]Published in https://pubmed.ncbi.nlm.nih.gov/27051591/

**Supp. Table 12.** Antibodies used for IP, WB and ICC

| | | | |
|---|---|---|---|
| α - α7nAChR<br>α - α7nAChR<br><br>α - D10<br>α - ULK4<br>α - Acetylated tubulin<br>α - α-tubulin<br><br>α – De-tyrosinated α-tubulin<br>α - βIII-tubulin<br><br>α - GAPDH<br><br>α - Histone H3<br><br><br>Alexa Fluor® 594 AffiniPure<br>Alexa Fluor® 488 AffiniPure | Mouse<br>Rat<br><br>Rabbit<br>Rabbit<br>Mouse<br><br>Mouse<br><br>Rabbit<br><br>Rabbit<br><br><br>Rabbit<br><br>Rabbit<br><br>Donkey anti-Rabbit IgG<br>Donkey Anti-Mouse IgG | Millipore-Sigma, Cat # M220<br>Santa Cruz Biotechnology Cat# sc-58607,<br>RRID: AB_784835<br>Abcam, Cat# ab198719<br>Fabgennix, Cat# ULK4-401AP<br>Sigma-Aldrich Cat# T7451, RRID: AB_609894<br><br>Cell Signaling Technology Cat# 3873, RRID: AB_1904178<br>Abcam, Cat# ab48389, RRID: AB_869990)<br><br>Cell Signaling Technology Cat# 5568, RRID: AB_10694505<br>Cell Signaling Technology Cat# 5174, RRID: AB_10622025<br>Cell Signaling Technology Cat# 4499, RRID: AB_10544537<br><br>Jackson ImmunoResearch Labs Cat# 711-585-152,<br>RRID: AB_2340621<br>Jackson ImmunoResearch Labs Cat# 715-545-151,<br>RRID: AB_2341099 | 1:2000 (WB)<br>IP, 1:500 (WB)<br><br>IP, WB (1:1000)<br>1:500 (WB), 1:100 (ICC)<br>1:3000(WB),<br>1:1000 (ICC)<br><br>1:1000 (WB),<br>1:200 (ICC)<br><br>1:1000 (WB)<br><br>1:200 (ICC)<br><br>1:1000 (WB)<br><br>1:1000 (WB)<br><br><br>1:500<br><br><br>1:500 |

### Supplementary Methods

#### RNA Isolation and RT-PCR

Total RNA was extracted from human brain tissue using Trizol (Invitrogen). Briefly, 20–30 mg of tissue was homogenized in Trizol with Lysing Matrix D (MP Biomedicals) by repeated vortexing, and RNA was purified following the manufacturer's protocol. cDNA was synthesized from 1 µg of total RNA using ImProm-II reverse transcriptase (Promega) and oligo (DT) primers at 42 °C for 1 h.

#### RNA-Seq Gene expression[42]

Bulk RNA sequencing of dorsolateral prefrontal cortex tissue from 205 NCI subjects from the ROSMAP study were utilized for *ULK4* gene expression analysis [43] . Genotype distribution, age, sex and PMI are depicted in Supplementary Table 1.
